# A novel terpene synthase produces an anti-aphrodisiac pheromone in the butterfly *Heliconius melpomene*

**DOI:** 10.1101/779678

**Authors:** Kathy Darragh, Anna Orteu, Kelsey J. R. P. Byers, Daiane Szczerbowski, Ian A. Warren, Pasi Rastas, Ana L. Pinharanda, John W. Davey, Sylvia Fernanda Garza, Diana Abondano Almeida, Richard M. Merrill, W. Owen McMillan, Stefan Schulz, Chris D. Jiggins

## Abstract

Terpenes, a group of structurally diverse compounds, are the biggest class of secondary metabolites. While the biosynthesis of terpenes by enzymes known as terpene synthases (TPSs) has been described in plants and microorganisms, few TPSs have been identified in insects, despite the presence of terpenes in multiple insect species. Indeed, in many insect species, it remains unclear whether terpenes are sequestered from plants or biosynthesised *de novo*. No homologs of plant TPSs have been found in insect genomes, though insect TPSs with an independent evolutionary origin have been found in Hemiptera and Coleoptera. In the butterfly *Heliconius melpomene*, the monoterpene (*E*)-β-ocimene acts as an anti-aphrodisiac pheromone, where it is transferred during mating from males to females to avoid re-mating by deterring males. To date only one insect monoterpene synthase has been described, in *Ips pini* (Coleoptera), and is a multifunctional TPS and isoprenyl diphosphate synthase (IDS). Here, we combine linkage mapping and expression studies to identify candidate genes involved in the biosynthesis of (*E*)-β-ocimene. We confirm that *H. melpomene* has two enzymes that exhibit TPS activity, and one of these, HMEL037106g1 is able to synthesise (*E*)-β-ocimene *in vitro*. Unlike the enzyme in *Ips pini*, these enzymes only exhibit residual IDS activity, suggesting they are more specialised TPSs, akin to those found in plants. Phylogenetic analysis shows that these enzymes are unrelated to previously described plant and insect TPSs. The distinct evolutionary origin of TPSs in Lepidoptera suggests that they have evolved multiple times in insects.

**Significance statement:** Terpenes are a diverse class of natural compounds, used by both plants and animals for a variety of functions, including chemical communication. In insects it is often unclear whether they are synthesised *de novo* or sequestered from plants. Some plants and insects have converged to use the same compounds. For instance, (*E*)-β-ocimene is a common component of floral scent and is also used by the butterfly *Heliconius melpomene* as an anti-aphrodisiac pheromone. We describe two novel terpene synthases, one of which synthesises (*E*)-β-ocimene in *H. melpomene*, unrelated not only to plant enzymes but also other recently identified insect terpene synthases. This provides the first evidence that the ability to synthesise terpenes has arisen multiple times independently within the insects.

## Introduction

Plants and insects sometimes use the same compounds for communication (1, 2). This may be adaptive if these chemicals exploit pre-existing sensory traits in the intended receiver. For example, sexually deceptive orchids mimic the scent of females of the pollinator species to attract males for pollination (3). Similarly, insects may use plant-like volatiles to exploit the sensory systems of other insects whose sensory systems have evolved for plant-finding (2, 4, 5). Phenotypic convergences such as these may involve different molecular mechanisms, including independent evolution at different loci, or may be due to the exchange of genes through horizontal gene transfer (6), and the concept has been studied across a range of organisms and phenotypes. However, we know little about the genetic basis of convergence in chemical signals.

One example of chemical convergence between plants and insects is the use of β-ocimene, a very common plant volatile, important in pollinator attraction due to its abundance and ubiquity in floral scents (7). This compound is also found in the genitals of male *Heliconius* butterflies (8–10). In *Heliconius melpomene*, (*E*)-β-ocimene acts as an anti-aphrodisiac pheromone, transferred from males to females during mating to repel further courtship from subsequent males (8). β-Ocimene is also found in large amounts in the flowers on which adult *H. melpomene* feed, and elicits a strong antennal response in both males and females (11, 12). This compound, therefore, appears to be carrying out two context-dependent functions, attraction to plants and repulsion from mated females.

Although β-ocimene synthases have been described in plants, none have been found in animals (7). It has previously been shown that *H. melpomene* is able to synthesise (*E*)-β-ocimene *de novo* (8). β-Ocimene is a monoterpene, a member of the largest and most structurally diverse class of natural products, the terpenes (13). Terpenes are formed from two precursors, isopentenyl diphosphate (IPP) and dimethylallyl diphosphate (DMAPP), which are themselves produced by either the universal mevalonate pathway or the methylerythritol phosphate pathway, the latter of which is absent in animals (14, 15). Varying numbers of IPP units are then added to DMAPP to form isoprenyl diphosphates of different chain lengths by isoprenyl diphosphate synthases (IDSs) (16, 17) (Fig. 1). These isoprenyl diphosphates are the precursors for the production of terpenes by terpene synthases (TPSs), with the length of the isoprenyl diphosphate determining the type of terpene that is made (18, 19). For example, DMAPP and one unit of IPP are converted by the IDS geranyl pyrophosphate synthase (GPPS) to form geranyl pyrophosphate (GPP), which can be converted by TPSs to monoterpenes, such as β-ocimene (Fig. 1). DMAPP and two units of IPP are converted by the IDS farnesyl diphosphate synthase (FPPS) into farnesyl diphosphate (FPP), which can be converted by TPSs to sesquiterpenes (20) (Fig. 1). Finally, DMAPP and three units of IPP are converted by the IDS geranylgeranyl diphosphate synthase (GGPPS) into geranylgeranyl diphosphate (GGPP), which can be converted by TPSs to diterpenes (Fig. 1).

**Figure 1:**
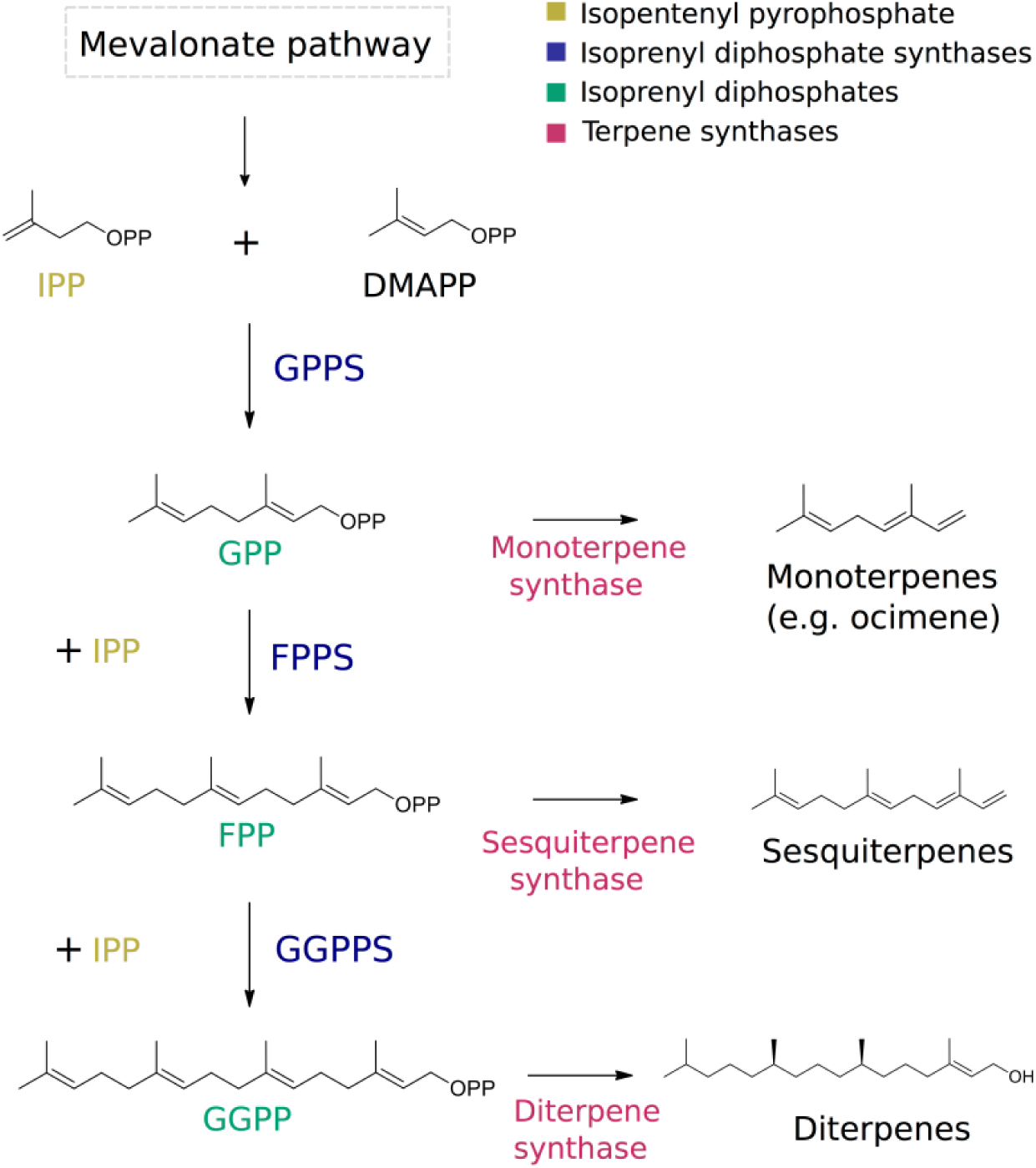
Pathway of terpene biosynthesis. Isopentenyl diphosphate (IPP) and dimethylallyl diphosphate (DMAPP) are first formed from the mevalonate pathway. IPP and DMAPP are the substrates for isoprenyl disphosphate synthases which produce isoprenyl diphosphates of varying lengths, depending on the number of IPP units added. Isoprenyl diphosphates are themselves the substrates used by terpene synthases to make terpenes of various sizes, for example, monoterpene synthases produce monoterpenes, such as ocimene, from geranyl pyrophosphate (GPP). For illustration, (E,E)-α-farnesene is used as a representative sesquiterpene, and phytol as a diterpene.

Both the mevalonate pathway, which forms IPP and DMAPP, and IDSs are ubiquitous in nature. In insects, the production of juvenile hormones is reliant on this pathway via FPP (21). In contrast, TPSs are more limited in their distribution. Until recently they had only been described in plants and fungi in the eukaryotic domain, suggesting that insects sequestered terpenes from their diet and were unable to synthesise these compounds *de novo* (15). In the last decade, insect TPS genes, which are not homologous to plant TPSs, have been discovered in Hemiptera and Coleoptera, and were shown to be involved in the production of aggregation and sex pheromones (1, 22–26). The enzymes found in Hemiptera are involved in the production of pheromone precursor sesquiterpenes from FPP, although the enzymes catalysing the terminal pheromone biosynthesis steps are unknown (25, 26). Sesquiterpene synthases have also been described in *Phyllotreta striolata* (Coleoptera) (24). The only monoterpene synthase described to date is that of *Ips pini* (Coleoptera), which produces a pheromone precursor from GPP (22, 23). These TPS genes have evolved from IDS-like genes, most closely related to FPPSs (1, 24). The TPS of *Ips pini* also retains IDS function, acting as both a GPPS and TPS *in vitro*. It is unclear whether the evolution of TPS activity occurred only once in insects, as the most recent phylogenetic evidence suggests, or has occurred independently in different lineages (1, 26).

Here, we identify the genes involved in the biosynthesis of (*E*)-β-ocimene in the butterfly *Heliconius melpomene* and analyse the evolution of terpene synthesis in *Heliconius* and other insects. To determine candidate TPS genes, we identified pathway orthologs in *H. melpomene* and carried out a genetic mapping study between *H. melpomene* and *H. cydno*, a closely-related species that does not produce (*E*)-β-ocimene. We identified a genomic region associated with the production of (*E*)-β-ocimene and searched for candidates within this region. We then identified genes with upregulated expression in the genitals of male *H. melpomene*, where (*E*)-β-ocimene is produced. We confirmed the TPS function of our candidate genes by expression in *E. coli* followed by enzymatic assays.

## Results

### *Expansion of IDSs in genome of* H. melpomene

We identified candidates potentially involved in terpene synthesis by searching in the genome of *H. melpomene* for enzymes in the mevalonate pathway and isoprenyl diphosphate synthases (IDSs) using well-annotated *Drosophila melanogaster* orthologs (Table S1)(21, 27, 28). We identified reciprocal best blast hits for all enzymes, except for acetoacetyl-CoA thiolase (Fig. 2). There was a clear one to one relationship for all enzymes, except for the IDSs which showed evidence for gene duplication. Of these, *Heliconius* contains two putatuive farnesyl diphosphate synthases (FPPSs), four putative copies of decaprenyl pyrophosphate synthase (DPPS) subunit two, and six putative geranylgeranyl pyrophosphate synthases (GGPPSs) (Fig. 2).

**Figure 2:**
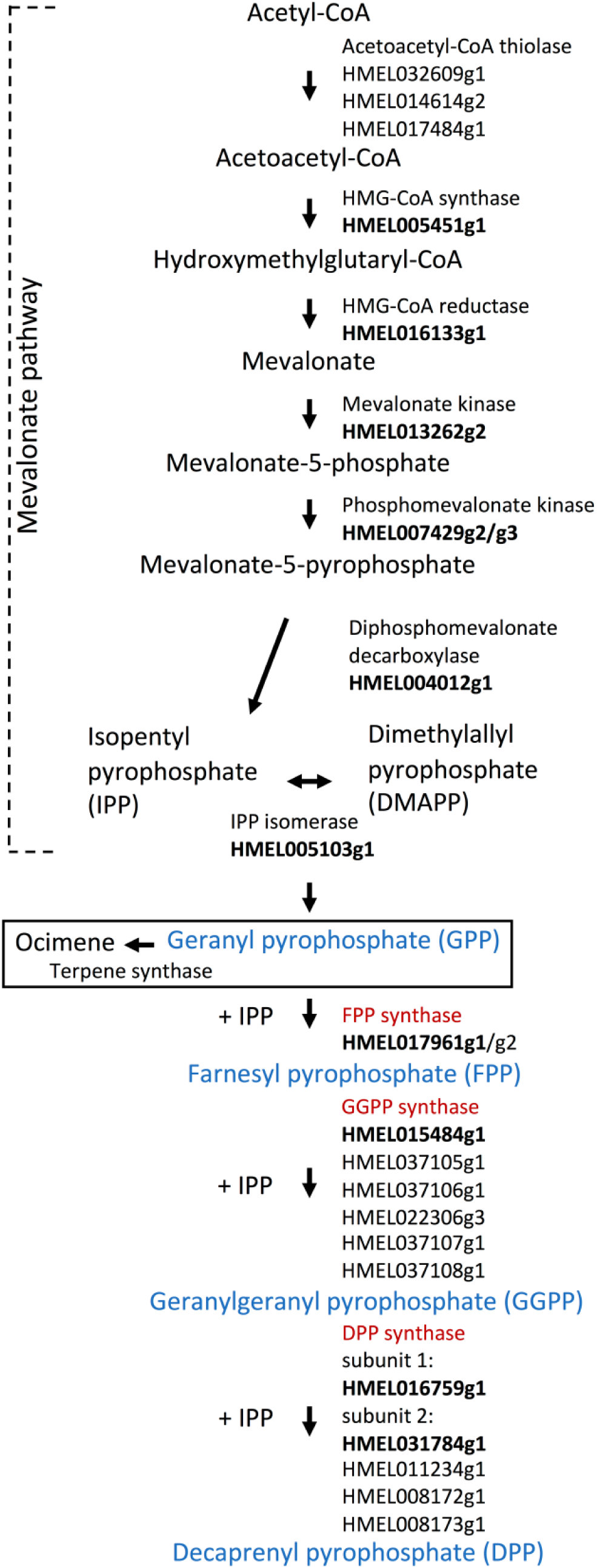
Proposed biosynthetic pathway in H. melpomene. Reciprocal best blast hits are highlighted in bold. IDSs are in red and their products, IDs, in blue. The first two exons of HMEL007429g2 and the last exon of HMEL007429g3 are expressed as a single transcript (for transcript sequence see Table S1).

The biggest expansion found was that of the GGPPSs, which are IDSs that catalyse the addition of IPP to FPP to form GGPP. One of these, *HMEL015484g1*, shows 83% amino acid sequence similarity to the GGPPS of the moth *Choristoneura fumiferana*, which has previously been characterised *in vitro* to catalyse the production of GGPP from FPP and IPP (29). *HMEL015484g1* is also the best reciprocal blast hit with the GGPPS of *D. melanogaster* (Fig. 2). The other five annotated GGPPSs show less than 50% similarity to the moth GGPPS, such that their function is less clear.

### QTL for (E)-β-ocimene production on chromosome 6

In order to determine which of the genes identified above could be important for (*E*)-β-ocimene production in *H. melpomene* we took advantage of the fact that a closely related species, *H. cydno*, does not produce (*E*)-β-ocimene (Fig. 3A). These two species can hybridise and, although the F1 females are sterile, F1 males can be used to generate backcross hybrids. We bred interspecific F1 hybrid males and backcrossed these with virgin females of both species to generate a set of backcross mapping families. The (*E*)-β-ocimene phenotype segregated in families backcrossed to *H. cydno* and so we focused on these families (Figure S1). Using quantitative trait locus (QTL) mapping with 114 individuals we detected a single significant peak on chromosome six associated with (*E*)-β-ocimene quantity (Fig. 3B). The QTL peak was at 36.4 cM, and the associated confidence interval spans 16.7-45.5 cM, corresponding to a 6.89Mb region containing hundreds of genes. The percentage of phenotypic variance explained by the peak marker is 16.4%, suggesting additional loci and/or environmental factor also contribute to the phenotype (Figure S2).

**Figure 3:**
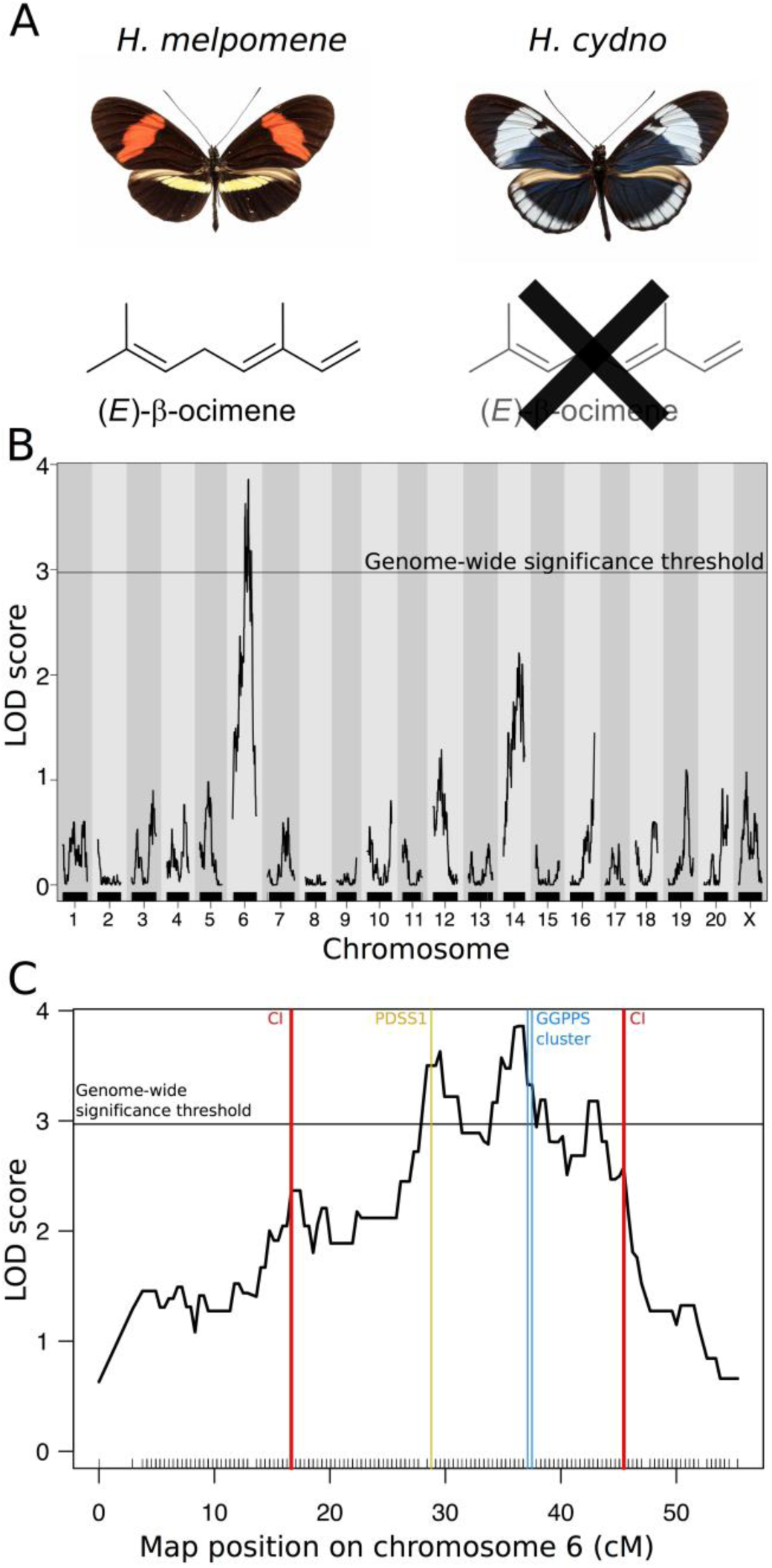
QTL for (E)-β-ocimene production. A) The two species used in the crosses, H. melpomene which produces (E)-β-ocimene, and H. cydno which does not. B) Genome-wide scan for QTL underlying (E)-β-ocimene production. C) QTL on chromosome 6 for (E)-β-ocimene production. Confidence intervals (CI) as well as the positions of candidate genes (subunit 1 of decaprenyl diphosphate synthase (PDSS1) and the GGPPS cluster) in the region are marked. Black lines above x-axis represent the position of genetic markers and horizontal line shows genome wide significance threshold (alpha=0.05, LOD=2.97).

### *Patterns of gene expression identify* HMEL037106g1 *and* HMEL037108g1 *as candidates*

To identify candidate genes for (*E*)-β-ocimene production we searched within the confidence interval of the QTL peak. We found that subunit 1 of DPPS, as well as all six GGPPSs were found in this region (Fig. 3C). We then compared the expression levels of the seven genes found within the QTL using published RNA sequencing (RNA-seq) data (30). We first analysed data from *H. melpomene* male and female abdomens and heads, mapped to the *H. melpomene* reference genome. Since (*E*)-β-ocimene is found in male abdomens in *H. melpomene*, we hypothesised that its synthase would be highly expressed in this sex and tissue. Only one gene showed male abdomen-biased expression: *HMEL037106g1* (Fig. 4A, Table S3, sex*tissue, t=-4.35, p<0.01). All other genes did not show a significant bias in this direction (Fig.4A, Table S3).

**Figure 4:**
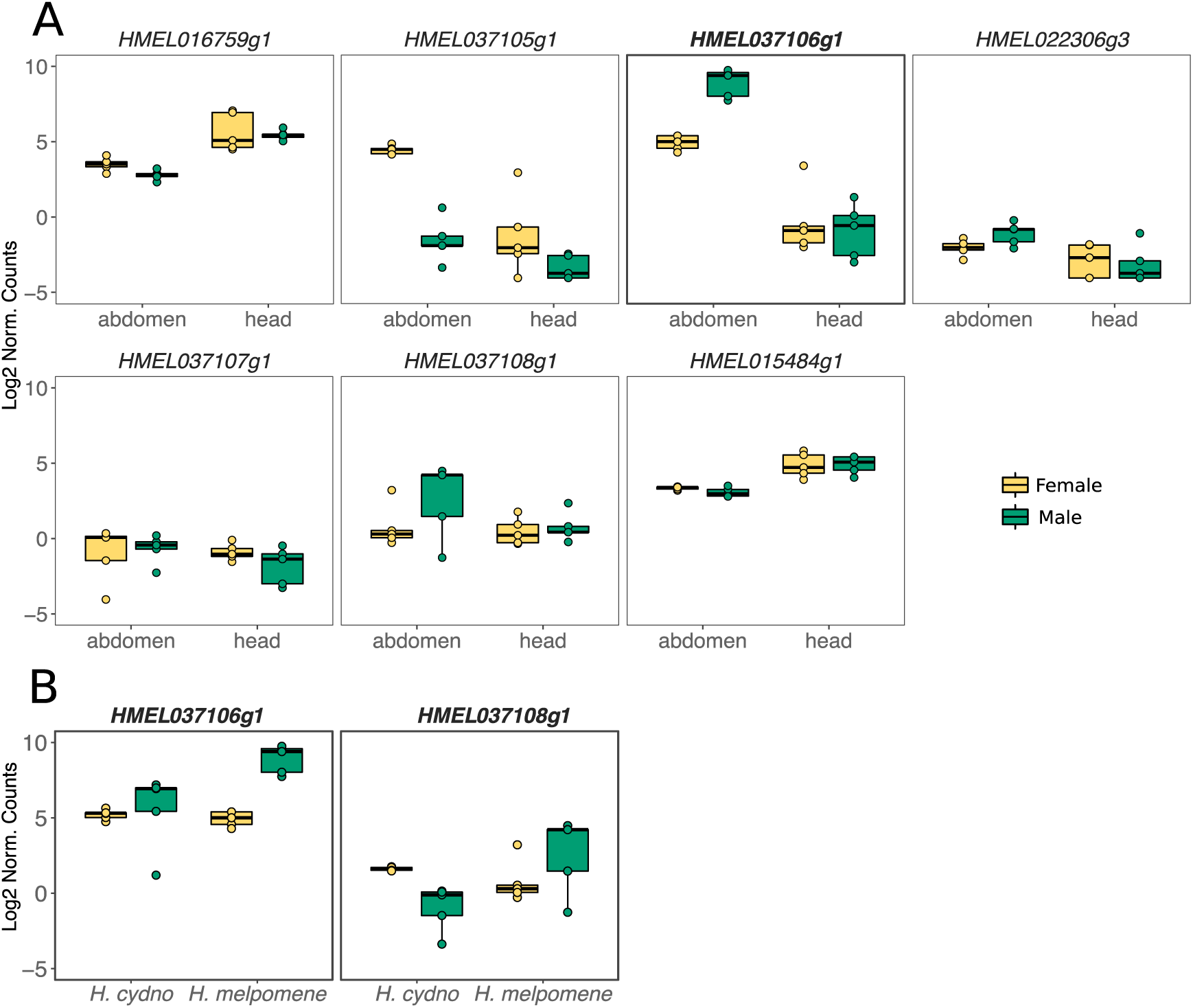
Gene expression analysis of candidate genes. A) Expression of genes in H. melpomene heads and abdomens of males and females. HMEL037106g1 (highlighted) shows male abdomen-biased expression. B) Expression of genes in H. melpomene and H. cydno abdomens of males and females (expression of other genes in Fig. S3). Both HMEL037106g1 and HMEL037108g1 (highlighted) show greater male-biased expression in H. melpomene than H. cydno. Full model statistics in Table S3 and S4. N=5 for each boxplot. Gene expression is given in log2 of normalised counts (using the TMM (trimmed mean of M values) transformation).

We next compared gene expression between *H. cydno* and *H. melpomene* abdomens. If HMEL037106g1 is synthesising (*E*)-β-ocimene, we expect its expression to be higher in *H. melpomene* male abdomens than in *H. cydno*, given that *H. cydno* does not produce the compound. We generated a reference-guided assembly of *H. cydno* by aligning an existing *H. cydno* Illumina trio assembly (31), to the *H. melpomene* reference, followed by automated gene annotation (see SI Materials and Methods). We then manually identified *H. cydno* orthologs for our seven candidate genes and checked for differential expression between species and sexes. *HMEL037106g1* and *HMEL037108g1* were the only genes showing greater male-biased expression in *H. melpomene* abdomens than in *H. cydno* abdomens (Fig. 4B, Table S4, *HMEL037106g1*, species*sex, t=3.15, p=<0.05; *HMEL037108g1*, species*sex, t=3.44, p<0.05). No other genes showed a significant bias in this direction (Fig. S3, Table S4). In summary, *HMEL037106g1* and to a lesser extent *HMEL037108g1* are primary candidate genes from within the QTL region.

### Functional experiments demonstrate the TPS activity of HMEL037106g1 and HMEL037108g1

We cloned HMEL037106g1 and HMEL037108g1 into plasmids and were able to generate heterologous expression of both proteins in *Escherichia coli*. We then conducted enzymatic assays with the expressed proteins using precursors from different points in the pathway to characterise their enzymatic function (Fig. 1, 5).

**Figure 5:**
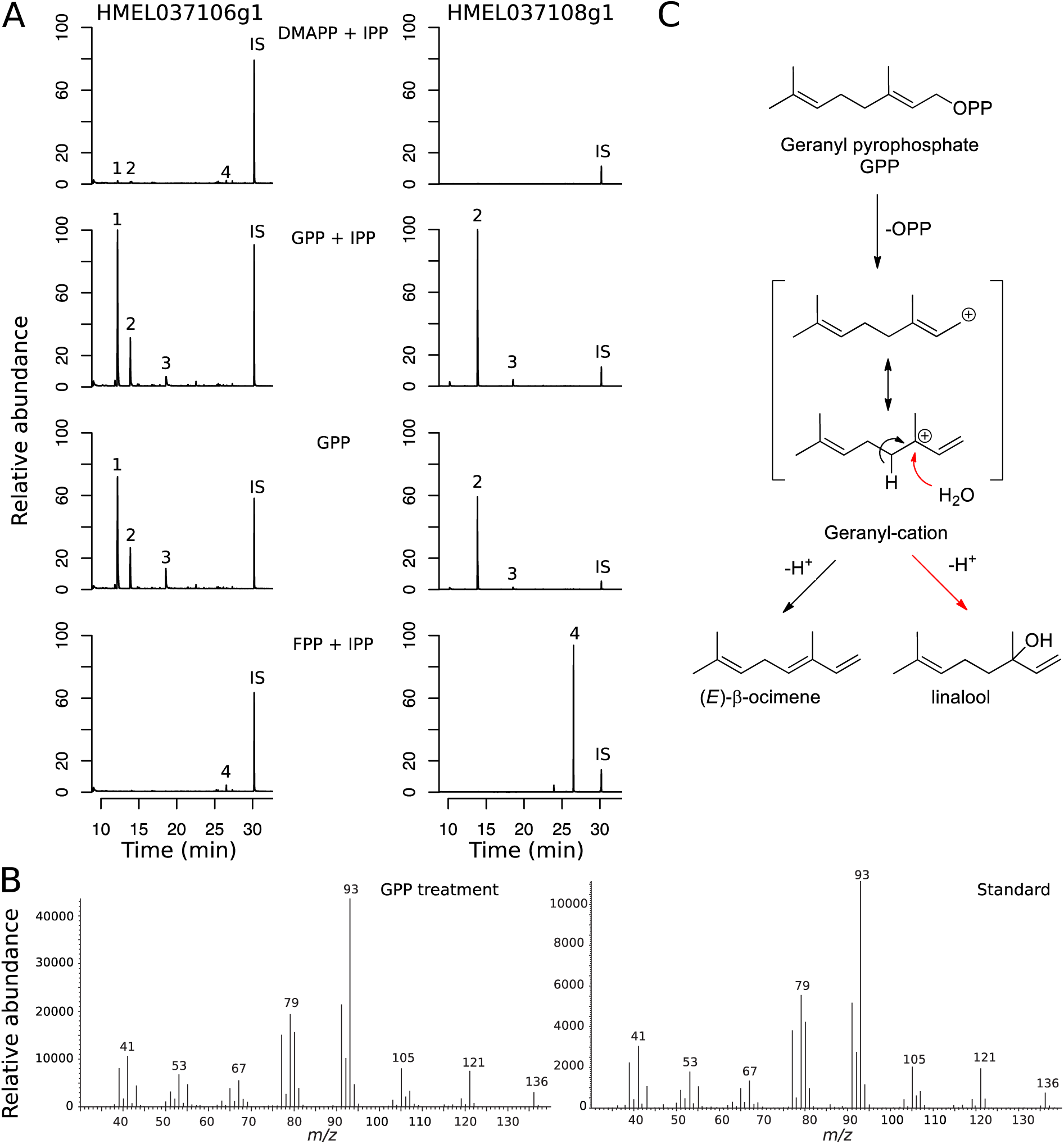
Functional characterisation of TPS activity of HMEL037106g1 and HMEL037108g1 from H. melpomene. A) Total ion chromatograms of enzyme products in the presence of different precursor compounds. HMEL037106g1 produces high amounts of (E)-β-ocimene in the presence of GPP, with trace amounts found in the treatment with DMAPP + IPP, and none with FPP. HMEL037108g1 produces large amounts of linalool with GPP, and nerolidol with FPP. 1, (E)-β-Ocimene; 2, Linalool; 3, Geraniol; 4, Nerolidol; IS, internal standard. Abundance is scaled to the highest peak of all treatments per enzyme. Quantification of peaks in Table S6 and S7. B) Confirmation of identity of (E)-β-ocimene by comparison of mass spectra of (E)-β-ocimene produced in experiments and a standard. C) Pathway of how (E)-β-ocimene and linalool are formed from GPP.

Firstly, we carried out assays with DMAPP and IPP, the two building blocks at the beginning of the terpene synthesis pathway to test for both IDS and TPS activity, as was seen in *Ips pini* (Fig. 1). HMEL037106g1 produced trace amounts of (*E*)-β-ocimene, linalool, another monoterpene, and nerolidol, a sesquiterpene, in this assay. This presumably occurs via the production of GPP and FPP, therefore HMEL037106g1 exhibits residual GPS and FPPS activity, as well as monoterpene synthase and sesquiterpene synthase activity to convert the GPP and FPP to (*E*)-β-ocimene, linalool, and nerolidol (Fig. 5A, Table S6). HMEL037108g1 produced trace amounts of linalool (Fig 5C) and nerolidol from DMAPP and IPP. Again, this demonstrates residual GPS and FPPS activity to form the GPP and FPP, and then both monoterpene and sesquiterpene synthase activity to convert these to linalool and nerolidol (Fig. 5A, Table S7).

We then carried out assays with GPP and IPP, as well as GPP alone to test for monoterpene synthase activity (Fig. 1). HMEL037106g1 showed monoterpene synthase activity, producing (*E*)-β-ocimene when provided with either GPP and IPP, or GPP alone (Fig. 5A, Table S6). Small amounts of (*Z*)-β-ocimene were also produced in treatments where (*E*)-β-ocimene was produced in large quantities (Table S6). In contrast to HMEL037106g1, HMEL037108g1 only produced (*E*)-β-ocimene in very small amounts from GPP (Table S9). Instead, linalool was produced in large amounts from GPP, suggesting that this enzyme is also acting as a monoterpene synthase but is responsible for production of linalool rather than (*E*)-β-ocimene (Fig. 5A, Table S7). HMEL037106g1 also produced linalool, albeit in much smaller quantities (Fig. 5A, Table S6).

Finally, we carried out assays with FPP and IPP to test for sesquiterpene synthase activity (Fig. 1). Although HMEL037106g1 exhibited small amounts of sesquiterpene synthase activity through the trace production of nerolidol from DMAPP and IPP (Table S6), when provided with FPP alone, sesquiterpene synthase activity was not demonstrated, suggesting it is not the primary enzyme function (Fig. 5A, Table S6). In contrast, HMEL037108g1 did exhibit sesquiterpene synthase activity, producing large amounts of nerolidol when FPP was provided as a precursor (Fig. 5C, Table S7).

Due to the linalool detected in treatments where (*E*)-β-ocimene was produced by HMEL037106g1, we tested whether linalool could be a metabolic intermediate between GPP and (*E*)-β-ocimene. However, HMEL037106g1 did not produce (*E*)-β-ocimene from linalool (Fig. S5, Table S8). The two stereoisomers of linalool, (*S*)-linalool and (*R*)-linalool, have different olfactory properties. We confirmed the stereochemistry of linalool produced by both enzymes and found that whilst HMEL037106g1 produced mainly (*S*)-linalool, HMEL037108g1 produced a racemic mixture (Fig. S6).

In summary, HMEL037106g1 is a monoterpene synthase, catalysing the conversion of GPP to (*E*)-β-ocimene (Fig. 5C, Table S9). HMEL037108g1 is a bifunctional monoterpene and sesquiterpene synthase catalysing the conversion of GPP to linalool as well as FPP to nerolidol (Fig. 5C, Table S9).

### Functional experiments demonstrate the residual IDS activity of HMEL037106g1 and HMEL037108g1

While the production of terpenes can be tested by direct GC/MS analysis of the products of each experiment, this method will not detect isoprenyl diphosphates, potentially missing IDS activity if it is present. In order to test for IDS activity, we repeated the above experiments with DMAPP and IPP, GPP and IPP, and FPP and IPP, followed by treatment with alkaline phosphatase to hydrolyse the isoprenyl diphosphate products to their respective alcohols. These alcohols can then be detected by GC/MS analysis.

No further IDS activity was detected in either enzyme, apart from the residual IDS activity already determined above due to the trace amounts of terpenes produced from DMAPP and IPP. When either enzyme is provided with GPP, geraniol is produced, and when provided with FPP, large amounts of farnesol is produced, as expected from the dephosphorylation of the provided precursors, and this is seen in control conditions as well (Fig. S7, S8, Table S10, S11). As expected from the previous experiments, (*E*)-β-ocimene is also produced when HMEL037106g1 is provided with GPP, and linalool and nerolidol are produced when HMEL037108g1 is provided with GPP and FPP, respectively. Geranylgeraniol is not produced in any treatments, demonstrating that neither HMEL037106g1 nor HMEL037108g1 is a GGPPS, as suggested by their annotation (Fig. S7, S8, Table S10, S11). In summary, both HMEL037106g1 and HMEL037108g1 only exhibit residual IDS activity.

### *Evolutionary history of gene family containing* Heliconius *TPSs*

Lineage-specific expansions of gene families are often correlated with functional diversification and the origin of novel biological functions (32). We therefore carried out a phylogenetic analysis of GGPPS in Lepidoptera to investigate whether gene duplication could have played a role in the evolution of the TPSs HMEL037106g1 and HMEL037108g1. Orthologs of the *H. melpomene* GGPPSs were identified in *H. cydno, Heliconius erato, Bicyclus anynana, Danaus plexippus, Papilo polytes, Pieris napi, Manduca sexta, Bombyx mori* and *Plutella xylostella* (33). Expansions of the GGPPS group of enzymes can be seen in *Heliconius* and in *Bicyclus*, both groups in which terpenes form part of the pheromone blend (34) (Fig. S9).

To focus on the *Heliconius*-specific duplications, we made a phylogeny using the DNA sequence of transcripts from *H. melpomene, H. cydno* and *H. erato. Heliconius melpomene* and *H. cydno* belong to the same clade within *Heliconius*, with an estimated divergence time around 1.5 million years ago (35). *Heliconius erato* is more distantly related, belonging to a different *Heliconius* clade which diverged from the *H. melpomene*/*H. cydno* group around 10 million years ago (36). Whilst (*E*)-β-ocimene is not found in the genitals of *H. cydno*, it is found in the genitals of *H. erato*, at around one tenth the amount of *H. melpomene* (37). We hypothesised that duplications between the *H. melpomene* and *H. erato* clades may have resulted in subfunctionalisation and a more efficient *H. melpomene* enzyme facilitating increased (*E*)-β-ocimene production. We found that both losses and gene duplications have occurred between the *H. melpomene* and *H. erato* clades, whilst gene copy number is conserved between closely-related *H. melpomene* and *H. cydno* (Fig. S10). The exact orthology between the *H. erato* and *H. melpomene*/*H. cydno* genes is unclear, but what is clear is that *H. melpomene*/*H. cydno* have more genes in this family than *H. erato* (Fig. S10), and that both clades have more genes than the ancestral lepidopteran state of one copy.

We also found evidence for the formation of pseudogenes following gene duplication. The amino acid sequences from translations of two genes in *H. melpomene, HMEL22305g1* and *HMEL037104g1*, do not contain complete functional protein domains. This is also seen for *Herato0606.241* in *H. erato*. Furthermore, more recent pseudogene formation could be seen in the *H. cydno* ortholog of *HMEL22306g3*, which contained multiple stop codons, despite exhibiting transcription (Fig. S3).

In order to determine the number of evolutionary origins of insect and plant TPSs we carried out a broader phylogenetic analysis, including other known insect and plant IDS and TPS proteins. Similar to the other insect TPSs described, *Heliconius* TPSs are not found within the same clade as plant ocimene synthases, representing an independent origin of ocimene synthesis in *Heliconius* and plants. Furthermore, the *Heliconiu*s TPSs do not group with known insect TPS enzymes in Hemiptera and Coleoptera (Fig. 6). Instead, the *Heliconius* TPS enzymes group with GPP and GGPP synthases, rather than FPP synthases. The TPS enzymes of *Heliconius* are therefore of an independent evolutionary origin to other insect TPSs.

**Figure 6:**
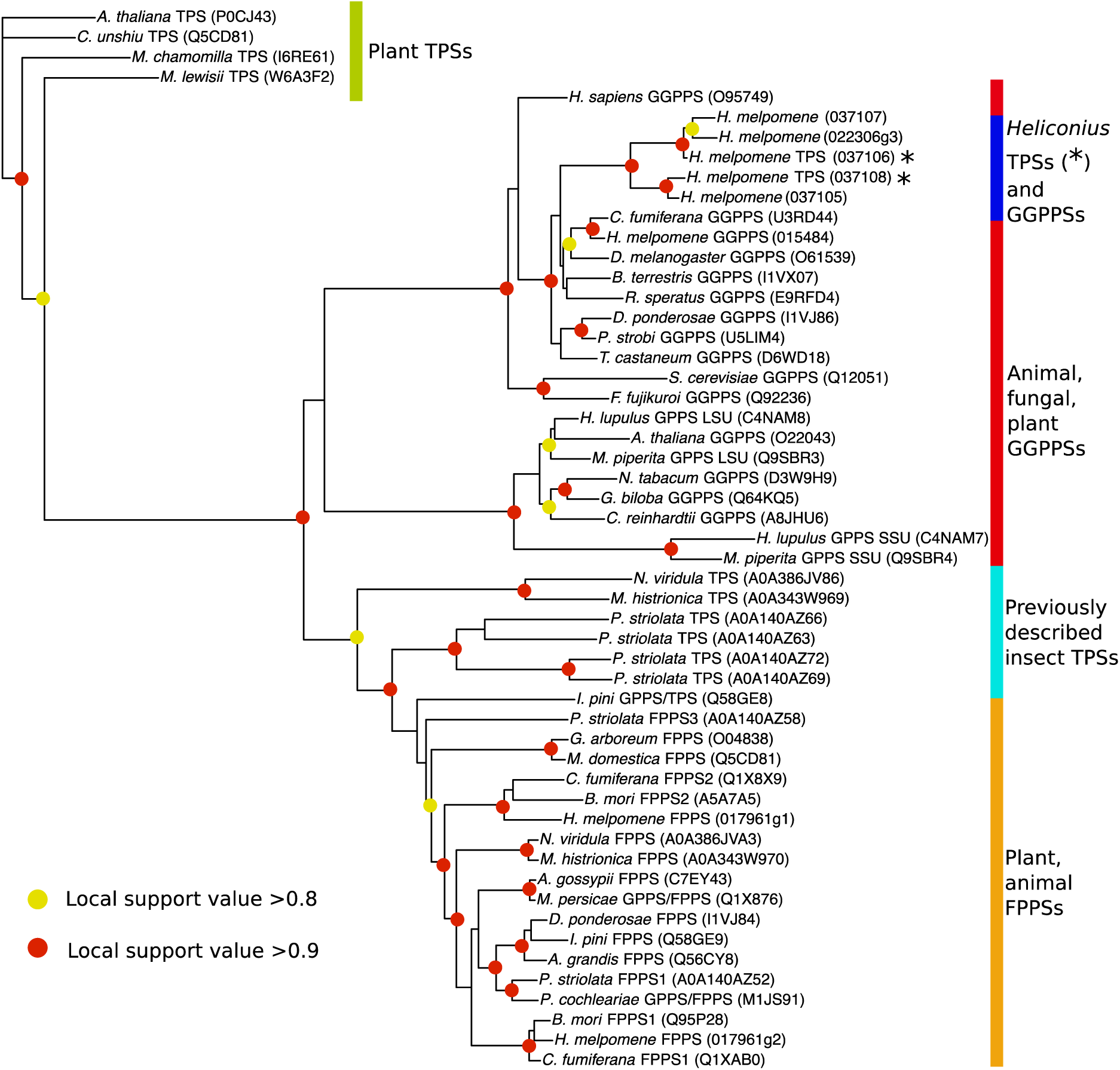
Phylogram of GGPPS, FPPS, and TPS proteins of animals, fungi, and plants. The phylogeny was constructed by FastTree using the JTT (Jones-Taylor-Thornton) model of amino acid evolution. Local support values are illustrated. The tree was rooted with the ocimene synthase of Citrus unshiu. Full species names in Table S12.

Comparison of the amino acid alignment of known insect TPSs with the *H. melpomene* enzymes (Fig. S11) demonstrated that residues previously identified as conserved in insect TPSs (25), were not found in the *H. melpomene* TPSs. No residues were shared between all insect TPSs (including *H. melpomene* TPS), which were not also shared with the *H. melpomene* GGPPS. This further indicates independent convergent evolution of TPS function in *H. melpomene*.

## Discussion

Both plants and animals use terpenes as chemical signals, however, the terpene synthases that make them have been identified in only a few insect species. Ocimene is a common monoterpene and we have identified the first ocimene synthase in animals. We identify a region of the genome responsible for differences in ocimene production and discover a novel gene family within this region. We confirm ocimene synthase activity for one of these enzymes (HMEL037106g1), and terpene synthase activity in a closely related enzyme (HMEL037108g1). Neither of these genes are homologous to known plant TPSs and represent a novel TPS family in *Heliconius*. Furthermore, they are also very different from previously described insect TPSs. While the TPS enzymes of Hemiptera and Coleoptera are more closely related to FPPSs (1, 24, 25), the *H. melpomene* TPSs are more closely related to GGPPSs. We do not find shared amino acid changes with other insect TPSs, strongly suggesting that TPS activity in Lepidoptera has arisen independently. Overall, the origin of the (*E*)-β-ocimene synthase activity in *H. melpomene* represents an excellent example of chemical convergence via the independent evolution of new gene function.

Male *Heliconius melpomene* transfer ocimene to the female during mating. Within this context the biological function of HMEL037106g1 is clear, making the anti-aphrodisiac compound (*E*)-β-ocimene. However, the *in vivo* role of HMEL037108g1 is less clear. It is found within the QTL interval and also shows higher expression in *H. melpomene* males relative to *H. cydno*. This enzyme acts as a multifunctional linalool/nerolidol synthase, which have previously been described in plants (38, 39). However, neither linalool or nerolidol are found in high amounts in the male abdomen (40). This apparent discrepancy may be due to the location or timing of expression *in vivo* (24). Another hypothesis is that *in vivo* GPP reacts with another substrate in the active site of HMEL037108g1, or that once linalool is produced it is metabolically channelled to another enzyme for further modification *in vivo* (41). This could also explain the lack of stereoselectivity in linalool formation. Further experiments will be required to determine if the other enzymes of this family not tested here exhibit TPS activity also.

Although we describe the first ocimene synthase in animals, ocimene synthases are likely to be found in other groups. Both *Bombus terrestris* and *Apis mellifera* use ocimene as a recruitment and larval pheromone, respectively (42, 43). While the biosynthetic pathway is not known in these groups, a similar pathway to that proposed here has been suggested in *A. mellifera* (44). However, the existing data suggest that the loci responsible for ocimene synthesis are also likely to be independently evolved. Unlike *H. melpomene*, only one GGPPS is found in the *Apis* genome, whilst there are six FPPS genes, the result of lineage-specific duplications (45). Although this needs to be confirmed by functional studies, based on the genomic patterns, we predict that convergence between Lepidoptera and Hymenoptera in the synthesis of ocimene also has an independent evolutionary origin.

Our findings also highlight the role that gene duplication plays in the evolution of new gene functions. Gene duplication is thought to be important for the evolution of new functions, as one gene copy can evolve a new function by a process called neofunctionalization (46), often resulting in large gene families with related but different functions. These families follow a birth-and-death model of evolution, expanding and contracting through gene duplication, formation of pseudogenes, and gene deletion (47, 48). Plant TPSs follow these dynamics, making up a large family formed of seven subfamilies, with lineage-specific expansions (49, 50). Gene duplication followed by neofunctionalization has resulted in closely-related enzymes which can produce different compounds, in some cases due to subcellular localisation (51).

Our data show a similar pattern of gene family diversification and suggest that gene duplication has facilitated the evolution of terpene synthesis in *Heliconius*. We uncover a lineage-specific expansion of GGPPSs in *Heliconius*. This novel gene expansion includes a number of pseudogenes, as well as two loci that possess TPS activity. A family of TPS genes has also been discovered in *Phyllotreta striolata*, where gene duplication is thought to have enabled functional diversification (24). Gene duplication can also facilitate enzyme specialisation by a process called subfunctionalization. In this case, an ancestrally multifunctional enzyme duplicates, resulting in two daughter copies which split the ancestral functions, and can result in optimisation of these two functions (46). Subfunctionalization might explain why neither *Heliconius* TPS shows significant IDS activity. In contrast to the multifunctional TPS/IDS enzyme from *I. pini* (22, 23), other insects have separate enzymes with IDS and TPS activity (24–26). One hypothesis is that an IDS enzyme initially gained TPS activity followed by gene duplication and subfunctionalization with enzymes specialised for different enzymatic steps. HMEL037106g1 is the first specialised monoterpene synthase described in animals.

We have identified a novel family of TPSs in *Heliconius* butterflies which is unrelated both to plant TPSs and to the few examples of previously described insect TPSs. We confirm that terpene synthesis has multiple independent origins in insects, which are themselves independent from the evolution of terpene synthesis in plants. Despite their independent evolution, insect TPSs show significant structural similarities, having evolved from IDS-like proteins. To understand how this diversity has arisen we need to identify the functional amino acid changes and relate structure to function, a nascent area of research for this group of enzymes (52).

## Materials and Methods

### *Analysis of biosynthetic pathway in* H. melpomene

To identify genes involved in terpene biosynthesis we searched the *H. melpomene* genome (v2.5) on LepBase (33, 53) for genes in the mevalonate pathway and IDSs. *Drosophila melanogaster* protein sequences were obtained from FlyBase and used in BLAST searches (blastp) against all annotated proteins in the *H. melpomene* genome (Table S1) (45, 54). We used the BLAST interface on LepBase with default parameters (-evalue 1.0e-10 -num_alignments 25) (33, 55). We then searched these candidate orthologs against the annotated proteins of *D. melanogaster* using the BLAST interface on FlyBase to identify reciprocal best blast hits. We included in our results other hits with an e-value smaller than 1e-^80^.

### Crossing for quantitative trait linkage mapping

To map the genetic basis of ocimene production we crossed *H. melpomene*, which produces (*E*)-β-ocimene, to *H. cydno*, a closely related species which does not. We crossed these two species to produce F1 offspring and backcross hybrids in both directions. Female F1s are sterile and so we mated male F1s to *H. cydno* and *H. melpomene* virgin stock females to create backcross families. Families created by backcrossing to *H. melpomene* had a phenotype similar to pure *H. melpomene*, suggesting the *H. melpomene* phenotype is dominant. While we used 265 individuals to create the linkage map, we focused on backcross families in the direction of *H. cydno*, where the (*E*)-β-ocimene phenotype segregates for the QTL mapping (Figure S1). We phenotyped and genotyped 114 individuals from 15 backcross families in the direction of *H. cydno*. Bodies were stored in dimethyl sulfoxide (DMSO) and stored at −20°C for later DNA extraction.

### Genotyping and linkage map construction

DNA extraction was carried out using Qiagen DNeasy kits (Qiagen). As previously described, individuals were genotyped either by RAD-sequencing (56–58), or low-coverage whole genome sequencing using Nextera-based libraries (57, 59). A secondary purification using magnetic SpeedBeads™ (Sigma) was performed prior to Nextera-based library preparation. Libraries were prepared following a method based on Nextera DNA Library Prep (Illumina, Inc.) with purified Tn5 transposase (59). PCR extension with an i7-index primer (N701–N783) and the N501 i5-index primer was performed to barcode the samples. Library purification and size selection was done using the same beads as above. Pooled libraries were sequenced by BGI (China) using HiSeq X Ten (Illumina).

Linkage mapping was conducted following (Byers et al., 2019), using standard Lep-MAP3(LM3) pipeline (60). Briefly, fastq files were mapped to the *H. melpomene* reference genome using BWA MEM (61). Sorted bams were then created using SAMtools and genotype likelihoods constructed (62). The pedigree of individuals was checked and corrected using IBD (identity-by-descent) and the sex checked using coverage on the Z chromosomes by SAMtools depth. A random subset of 25% of markers were used for subsequent steps. Linkage groups and marker orders were constructed based on the *H. melpomene* genome and checked with grandparental data.

The map constructed contained 447,820 markers. We reduced markers by a factor of five evenly across the genome resulting in 89,564 markers with no missing data to facilitate computation. We log-transformed amounts of ocimene produced to conform more closely to normality. Statistical analysis was carried out using R/qtl (Broman et al. 2003). We carried out standard interval mapping using the *scanone* function with a non-parametric model, an extension of the Kruskal-Wallis test statistic. The analysis method for this model is similar to Haley-Knott regression (Haley and Knott 1992). We used permutation testing with 1000 permutations to determine the genome-wide LOD significance threshold. To obtain confidence intervals for QTL peaks we used the function *bayesint.* Phenotype data, pedigree, linkage map and R script is available from OSF (https://osf.io/3z9tg/?view_only=63ba7c0767a84d8eb907fbf599df062f). Sequencing data used for the linkage maps is available from ENA project ERP018627 (57). GC/MS data is available from Dryad Data Repository (XXXX).

### *Phenotyping of (*E*)-β-ocimene production*

Chemical extractions were carried out on genital tissue of mature (7-14 days post-eclosion) male individuals of *H. melpomene, H. cydno*, and hybrids (for details of butterfly stocks please see SI Material and Methods). Genitals were removed using forceps and soaked, immediately after dissection, in 200μl of dichloromethane containing 200 ng of 2-tetradecyl acetate (internal standard) in 2ml glass vials with polytetrafluoroethylene-coated caps (Agilent, Santa Clara, California). After one hour, the solvent was transferred to a new vial and stored at −20°C until analysis by gas chromatography-mass spectrometry (GC-MS). For details of GC-MS analysis please see SI Materials and Methods.

### RNA sequencing analysis

Gene expression analyses were performed using already published RNA-seq data from heads and abdomens of *H. melpomene* and *H. cydno* from GenBank BioProject PRJNA283415 (30). Although it would be possible to map the H.cydno RNA-seq reads to H. melpomene due to high genome sequence similarity, that might lead to biases associated with reads carrying H. cydno specific alleles. To accurately quantify gene expression in H. cydno we generated an assembly and annotation of the H. cydno genome using sequencing data available from ENA study ERP009507 (56). We then manually identified the *H. cydno* orthologs of our seven candidate genes (Table S2) and curated the annotation to make it compatible with RNA-seq analysis software (for details of the assembly and annotation please see SI Materials and Methods). We performed quality control and low quality base and adapter trimming on the RNA-seq data using TrimGalore! (63). We then mapped the reads to the *H.melpomene* genome v2.5 (64) and our newly assembled *H.cydno* genome using STAR (65). *featureCounts* (66) was used to produce read counts that were normalised by library size with TMM (trimmed mean of M values) normalisation (67) using the edgeR package in R (68). To test for differences in expression of our candidate genes, we used the *voom* function from the limma package in R (69), which fits a linear model for each gene by modelling the mean-variance relationship with precision weights.

To test for male abdomen-biased expression within *H. melpomene* we included two fixed effects, sex and tissue, as well as including individual as a random effect (expression ∼ sex + tissue + sex*tissue + (1|individual)). We were looking for genes with a significant interaction between sex and tissue, showing higher expression in male abdomens. To test for differences in expression between *H. melpomene* and *H. cydno* abdomens we included two fixed effects, sex and species, as well as an interaction term (expression ∼ sex + species + species*tissue). We were interested in finding differences in the extent of sex-bias between species, again detected by a significant interaction term with higher expression in *H. melpomene* male abdomens.

P-values were corrected for multiple testing using the Benjamini-Hochberg procedure for all genes in the genome-wide count matrix (17902 for *H. melpomene*). For the interspecific comparison we identified genome wide orthologs from the annotation and produced a gene count matrix including both species. The ortholog list was limited to genes that had only one ortholog in each species (11571 genes). Scripts are available from OSF (https://osf.io/3z9tg/?view_only=63ba7c0767a84d8eb907fbf599df062f).

### In vitro *expression and enzymatic assays*

For more details of the expression and enzymatic assays please see SI Materials and Methods. Briefly, cDNA libraries were synthesised from RNA extracted from male abdominal tissue of *H. melpomene*. Genes of interest were amplified by PCR with gene-specific primers (Table S5), purified, sequenced for confirmation, ligated into expression plasmids, and transformed into competent *Escherichia coli* cells. Cell cultures were grown to an OD_600_ of 0.5 were induced with 1mM IPTG and cultivated for a further two hours before collection by centrifugation. Cells were resuspended in assay buffer and sonicated.

TPS and IDS activity was assayed using the soluble fraction of the cell lysate. We added either DMPP and IPP, GPP and IPP, FPP and IPP, or GPP alone. We also tested for enzymatic activity with (*R*)-linalool and (*S*)-linalool. For TPS activity assays, reactions were immediately stopped on ice and extracted with hexane. For IDS activity assays, reaction mixtures were incubated with alkaline phosphatase to hydrolyse the pyrophosphates before hexane extraction. Prior to analysis by GC-MS, an internal standard was added and samples concentrated. Products were compared to control experiments without protein expression. For details of GC-MS analysis please see SI Materials and Methods. GC/MS data is available from Dryad Data Repository (XXXX).

### Phylogenetic analysis

To identify orthologs of the GGPPS in other Lepidoptera we searched protein sequences from *H. melpomene* version 2.5 (56, 64) against the genomes of *H. erato demophoon* (v1), *Bicyclus anynana* (v1×2), *Danaus plexippus* (v3), *Papilo polytes* (ppol1), *Pieris napi* (pnv1×1), *Manduca sexta* (msex1), *Bombyx mori* (asm15162v1), and *Plutella xylostella* (pacbiov1), using the BLAST interface (tblastn) on LepBase (33, 55). We also included the previously identified orthologs from the *H. cydno* genome (Table S2). To check that the predicted orthologs contained functional protein domains we used the NCBI conserved domain search with default parameters (70). We deleted any proteins found without complete functional domains, including a gene from *H. erato, Herato0606.241*. We also did not include the *H. cydno* ortholog of *HMEL22306g3* in the protein tree, as despite showing transcription (Fig. S3), there were multiple stop codons within the coding region.

To focus on the *Heliconius*-specific duplications, we downloaded the transcript sequences for the *H. melpomene* and *H. erato* proteins from LepBase and exported transcripts for predicted genes in Apollo for *H. cydno*. (Table S2). We used gene *Herato0606.245* (GGPPS, shows high similarity to the GGPPS of the moth *Choristoneura fumiferana*) to root the tree.

To investigate the evolutionary relationship of the *Heliconius* GGPPS we carried out a broader phylogenetic analysis with other known insect and plant IDS and TPS proteins. Protein sequences for these additional enzymes were downloaded from Uniprot (71). *Heliconius* protein sequences were obtained as described above. We used an ocimene synthase enzyme from *Citrus unshiu* to root the tree.

We aligned amino acid or DNA sequences using Clustal Omega on the EMBL-EBI interface (72). Alignments were visualised using BoxShade (https://embnet.vital-it.ch/software/BOX_form.html). Phylogenies were inferred using FastTree, a tool for creating approximately-maximum-likelihood phylogenetic trees, with default parameters (73, 74). These phylogenies were plotted using the package *ape* and *evobiR* in R version 3.5.2. (75–77). To ensure correct placement of support values when re-rooting trees we checked phylogenies using Dendroscope (78, 79). Phylogenies and R code are available from OSF (https://osf.io/3z9tg/?view_only=63ba7c0767a84d8eb907fbf599df062f).

## Data availability

The *H. cydno* assembly is available from OSF (https://osf.io/3z9tg/?view_only=63ba7c0767a84d8eb907fbf599df062f) and was assembled using previously published sequencing data available from ENA study ERP009507 (56). Sequencing data used to make linkage maps is available from ENA study ERP018627 (57). RNA sequencing data of *H. cydno* and H. *melpomene* heads and abdomens was obtained from GenBank BioProject PRJNA283415 (30). Raw data and scripts used for analysis are available from OSF (https://osf.io/3z9tg/?view_only=63ba7c0767a84d8eb907fbf599df062f). GC/MS data is available from Dryad Data Repository (XXXX).

## Acknowledgements

We thank the team at the insectaries in Panama, including Oscar Paneso, for help rearing butterflies. We thank Marek Kučka and Yingguang Frank Chan for providing the Tn5 enzyme used for the preparation of sequencing libraries. KD and AO were supported by the Natural Research Council Doctoral Training Partnership (grant number (grant NE/L002507/1) and KD additionally by a Smithsonian Tropical Research Institute Short Term Fellowship. KJRPB, IAW, RMM and CDJ were supported by the European Research Council (grant 339873 SpeciationGenetics). RMM was also supported by a Deutsche Forschungsgemeinschaft Emmy Noether fellowship (grant GZ:ME4845/1-1). AP was supported by a Natural Research Council studentship (PFZE/063) and a Smithsonian Tropical Research Institute Short Term Fellowship. JWD was funded by a Herchel Smith Postdoctoral Research Fellowship and a Smithsonian Tropical Research Institute Fellowship. WOM was supported by the Smithsonian Tropical Research Institute and National Science Foundation (grant DEB 1257689). SS thanks the Deutsche Forschungsgemeinschaft (grant Schu984/12-1) supporting DS.

