## Supplementary file for "A novel terpene synthase produces an anti-aphrodisiac pheromone in the butterfly *Heliconius melpomene*"

PNAS
www.pnas.org

Supplementary Information for

### A novel terpene synthase produces an anti-aphrodisiac pheromone in *Heliconius melpomene*

Kathy Darragh, Anna Orteu, Kelsey J. R. P. Byers, Daiane Szczerbowski, Ian A. Warren, Pasi Rastas Ana L. Pinharanda, John W. Davey, Sylvia Fernanda Garza, Diana Abondano Almeida, Richard M. Merrill, W. Owen McMillan, Stefan Schulz, Chris D. Jiggins

Corresponding author: Kathy Darragh

**This PDF file includes:**

Supplementary text

References for supplementary text

Figs. S1 to S11

Tables S1 to S12

### SI Materials and Methods

#### *Butterfly stocks*

Outbred stocks of *Heliconius melpomene rosina* and *H. cydno chioneus* were established from wild individuals collected in Gamboa (9˚7.4’ N, 79˚42.2’ W, elevation 60 m) in the nearby Soberania National Park, San Lorenzo National Park (9°17’N, 79°58’W; elevation 130 m), and in Altos de Campana National Park (8°69’ N, 79°92’ W; elevation 900 m). Stocks were maintained in insectaries at the Smithsonian Tropical Research Institute (STRI) facilities in Gamboa, Panama. Individuals for this study were reared under ambient conditions between January 2016 and January 2018. Larvae were reared on *Passiflora platyloba*. Adult male butterflies were kept in cages with other males and provided with approximately 20% sucrose solution with access to at least one of *Psychotria poeppigiana*, *Gurania eriantha*, *Psiguiria triphylla* and *Psiguria warscewiczii* as pollen sources.

#### *GC/MS analysis*

DMAPP (90%), GPP (95%) and IPP (95%) were purchased from Sigma. FPP was purchased from VWR (98%). (*R*)-Linalool was purchased from Merck (95%). (*S*)-Linalool was purchased from Sigma as coriander oil and purified through column chromatography.

Extracts from adult butterflies, and samples from *in vitro* experiments were analysed by gas chromatography/mass spectrometry (GC/MS) using an Agilent (model 5977/5975) mass-selective detector connected to an Agilent GC (model 7890B/7890A) with electron impact ionisation (70eV). This instrument was equipped with an Agilent ALS 7693 autosampler and an HP-5MS fused silica capillary column (Agilent) (length 30 m, inner diameter 0.25 mm, film thickness 0.25 µm). Injection was performed in splitless mode (injector temperature 250°C) with helium as the carrier gas (constant flow of 1.2 ml/min). The temperature programme started at 50°C for 5 min, and then rose at a rate of 5°C /min to 320°C, before being held at 320°C for 5 min. The compounds were identified through comparison with retention time and mass spectra of standard samples.

Chiral analysis of Linalool was performed in an Agilent 7820A gas chromatograph equipped with flame ionization detector (FID), using a chiral column Beta DEX 225 (length

30m, inner diameter 0.25mm, film thickness 0.25µm, Supelco). The oven program started at 50°C for 1 minute, followed by increasing the temperature at a rate of 3°C/min until 210°C, keeping this temperature for 5 minutes. Samples were injected in splitless mode and flow of 1.65 mL/min. The peak areas were used to calculate the percentage of each stereoisomer.

##### Heliconius cydno *guided assembly and annotation transfer*

*H. cydno* and *H. melpomene* had their most recent common ancestor 1.5 million years ago and their absolute divergence is roughly 3% (dxy ~0.03) (1, 2). Due to this high degree of similarity, it is possible to map *H. cydno* RNA-seq reads to the *H. melpomene* genome. However, we wanted to accurately quantify gene expression in existing *H. cydno* samples (GenBank BioProject PRJNA283415 (3)) by reducing potential biases associated with RNA-seq reads carrying H. *cydno* specific alleles. RNA-seq reads from *H. cydno* with such variants have a lower probability to map correctly to the existing *H. melpomene* reference. This biases quantification and increases false positive rates, documented extensively, specifically in the context of allele-specific expression studies. (4).

A *H. cydno* trio (mother, father and progeny) was previously Illumina sequenced (ENA study ERP009507) and assembled into maternal and paternal genomes with trio-sga (5). The paternal genome had 34,566 scaffolds, a total size of 270,339,622 bp and a scaffold N50 of 25,716 bp, with 551 kb of gaps (paternal trio fasta file available from OSF (https://osf.io/3z9tg/?view_only=63ba7c0767a84d8eb907fbf599df062f)). To improve gene contiguity we used the progressiveCactus algorithm (v3) to align the *H. cydno* paternal haplotypic assembly to the chromosomal version of the *H. melpomene* genome (v2.5, 6–8). The HAL database created by progressiveCactus was loaded to Ragout (v1.2, 9) to produce the final *H. cydno* reference-guided assembly (*H. cydno* reference fasta file; ordering information and unplaced scaffolds available from OSF (https://osf.io/3z9tg/?view_only=63ba7c0767a84d8eb907fbf599df062f). The *H. cydno* guided assembly has 58 scaffolds, a total size of 261,056,210 bp, a scaffold N50 of 13,724,118 bp, and 8.3 Mb of gaps.

We then transferred the *H. melpomene* annotation (v2.5) to the *H. cydno* assembly. We used EMBOSS Seqret (v6.6.0.0) to convert the *H. melpomene* annotation file to the embl format (10) and we used RATT to transfer the *H. melpomene* annotation (reference) to the guided *H. cydno* genome (query). RATT is part of PAGIT, a post-assembly genome-improvement toolkit (v1.0) (11). We searched for synteny between the reference and the query using MUMmer (v4.0) and detected possible errors such as start and stop codons or frameshift mutations (12). After correcting such errors with the RATT pipeline the annotation transfer to *H. cydno* was complete (13).

To ensure that our genes of interest from *H. melpomene* (those identified in Table S1) were correctly annotated we manually curated these genes in the *H. cydno* annotation. To find orthologs in *H. cydno* we used the BLAT function in Apollo to search for *H. melpomene* exons (14, 15). We checked the gene models for splice sites and start and stop codons. The curated gene models were then exported from Apollo and manually included in the *H. cydno* annotation. We then subset the annotation to include only exons, because CDS sequences had not been properly annotated (Updatedannotation.gff). We then converted it to gtf file format using the gffread function of Cufflinks (Hcyd1.0_annotV2.gtf) (16) and filtered out exons longer than 30,000 base pairs (Hcyd1.0_annotV2.gtf; gtf_modify_Hcyd_annotV2.R). We finally used the gtf_modify_Hcyd_annotV3.R script to include unique *H. cydno* gene-ids (Hcyd1.0_annotV3.gtf).

##### In vitro *expression and enzymatic assays*

RNA extraction from male abdominal tissue of *H. melpomene* was carried out following a standard TRIzol protocol (Invitrogen) and cDNA synthesised using 5x iScript Reaction Mix (Bio-Rad). Following the protocol from *Champion™ pET101 Directional TOPO™ Expression Kit* (Invitrogen), we amplified the full-length transcript of genes of interest from the cDNA by PCR using Q5 High-Fidelity 2x Master Mix (Biolabs), with gene-specific primers (Table S5). The primers were designed for full-length transcript amplification. The PCR products were purified using a MiniElute PCR purification kit (Qiagen) and then sequenced to confirm identity (Table S5). Following sequencing, the PCR products were ligated into the expression vector pET101/D-TOPO® and transformed into One Shot® TOP10 Chemically Competent *Escherichia coli* cells. Plasmids were extracted from cultures of successful colonies using the QIAprep Spin Miniprep Kit (Qiagen) and sequenced again to confirm correct ligation in the vector using the T7 and T7-reverse primers (Table S5).

Plasmids containing the genes of interest in the correct orientation were transformed into *Escherichia coli* strain BL21 Star™(DE3) for expression. Cell cultures were grown to an $OD_{600}$ of 0.5 and induced with 1mM IPTG. After induction, the cells were cultivated for a further two hours at 37°C and 250rpm, before collection by centrifugation for 15 minutes at 6000xg at 4°C. Expression of protein was verified using sodium dodecyl sulfate polyacrylamide gel electrophoresis (SDS/PAGE). Pellets were resuspended in chilled extraction buffer (25 mM 4-(2-hydroxyethyl)-1-piperazineethanesulfonic acid pH 7.5, 1 mM MnCl2, 100 mM KCl, 3 mM dithiothreitol, 10% glycerol, protease inhibitor cocktail (Sigma)) and disrupted by sonication. Cell lysates were then centrifuged for 10 minutes at 9000xg at 4°C and the supernatant (containing the soluble part of the cell lysate) retained.

TPS and IDS activity was assayed using the soluble fraction of the cell lysate. Protein concentration was estimated using a Qubit® Protein Assay Kit (Invitrogen). 80-100ng of protein was added to each reaction in a total volume of 300µl. We added different precursors from different steps in the pathway (Fig. 1) to characterise enzymatic activity. Experiments were incubated at 30°C for two hours at 200rpm.

Firstly, we added DMAPP and IPP (100µM each), the two building blocks at the beginning of the pathway. To form a terpene from these compounds, they first need to be combined to form GPP, which can then be converted to a terpene by TPS activity. If the enzyme is a multifunctional GPPS/TPS, as in *Ips pini*, monoterpenes should be formed from DMAPP and IPP, via the production of GPP. Furthermore, if FPPS or GGPPS activity is present, FPP or GGPP could be formed from DMAPP and IPP, as well as sesquiterpene or diterpenes if sesquiterpene or diterpene synthase activity is exhibited. We then carried out assays with GPP (100µM) and IPP (50µM). If the enzyme solely exhibits monoterpene synthase activity, the monoterpene could only be formed from GPP directly and not from DMAPP and IPP. Furthermore, the enzyme could be an FPPS or GGPPS, and could therefore produce FPP or GGPP from GPP and IPP. FPP and GGPP could be converted to sesquiterpenes or diterpenes if sesquiterpene or diterpene synthase activity is exhibited. We also tested with GPP alone (100µM) to test for monoterpene synthase activity directly. Finally, we carried out assays with FPP (100µM) and IPP (50µM). If the enzyme is a GGPPS as annotated, it should form GGPP from FPP and IPP, as well as potentially converting GGPP to diterpenes. This is also a test for sesquiterpene synthase activity, as sesquiterpenes should be formed from FPP if the enzyme is a sesquiterpene synthase. We also tested for enzymatic activity with (*R*)-linalool and (*S*)-linalool (100µM).

To test for IDS activity, we repeated the above experiments with DMAPP and IPP, GPP and IPP, and FPP and IPP, followed by treatment with alkaline phosphatase to hydrolyse the isoprenyl diphosphate products to their respective alcohols. These alcohols can then be detected by GC/MS analysis.

Dephosphorylation of GPP produces the monoterpene alcohol geraniol, whilst dephosphorylation of FPP produces the sesquiterpene alcohol farnesol. Our expectations for controls, without IDS or TPS enzymatic activity, is to find geraniol when GPP is provided, and farnesol when FPP is provided. Linalool is a monoterpene alcohol which is an isomer of geraniol, and nerolidol is a sesquiterpene alcohol which is an isomer of farnesol. Furthermore, geranylgeraniol is a diterpene alcohol derived from the dephosphorylation of GGPP. If an enzyme is exhibiting IDS activity, we expect it to be able to catalyse the condensation of IPP and the other precursor provided, DMAPP, GPP, or FPP, to form larger molecules. Therefore, when provided with DMAPP and IPP, we expect to find either monoterpene or sesquiterpene alcohols, derived from GPP or FPP. When provided with GPP and IPP, we expect sesquiterpene alcohols derived from FPP. When provided with FPP and IPP, we expect larger diterpene alcohols, such as geranylgeraniol, to form via the formation of GGPP.

For TPS activity assays, reactions were stopped on ice and overlaid with 250µl hexane and left at 25°C overnight. The hexane layer was then transferred to a new vial and stored at -80°C. For IDS activity assays, following incubation with the precursors, 20 units of alkaline phosphatase (Sigma) in alkaline phosphatase buffer was added to each reaction mixture and incubated at 30°C for four hours at 200rpm. Following this, 250µl hexane was added and left at 25°C overnight. The hexane layer was then transferred to a new vial and stored at -80°C. Prior to analysis by GC/MS, 20 µL of a solution of 2-acetoxytetradecane in hexane (10 µg/mL) was added as an internal standard, and samples concentrated to a volume of approximately 30 µL. Products were compared to control experiments without protein expression.

***Figure S1:*** *Amount of* (E)-β-*ocimene (ng) in both pure parental species, F1 hybrids, and backcrosses in both directions. The phenotype segregates in backcrosses to* H. cydno *and, therefore, we focused on this cross direction.*

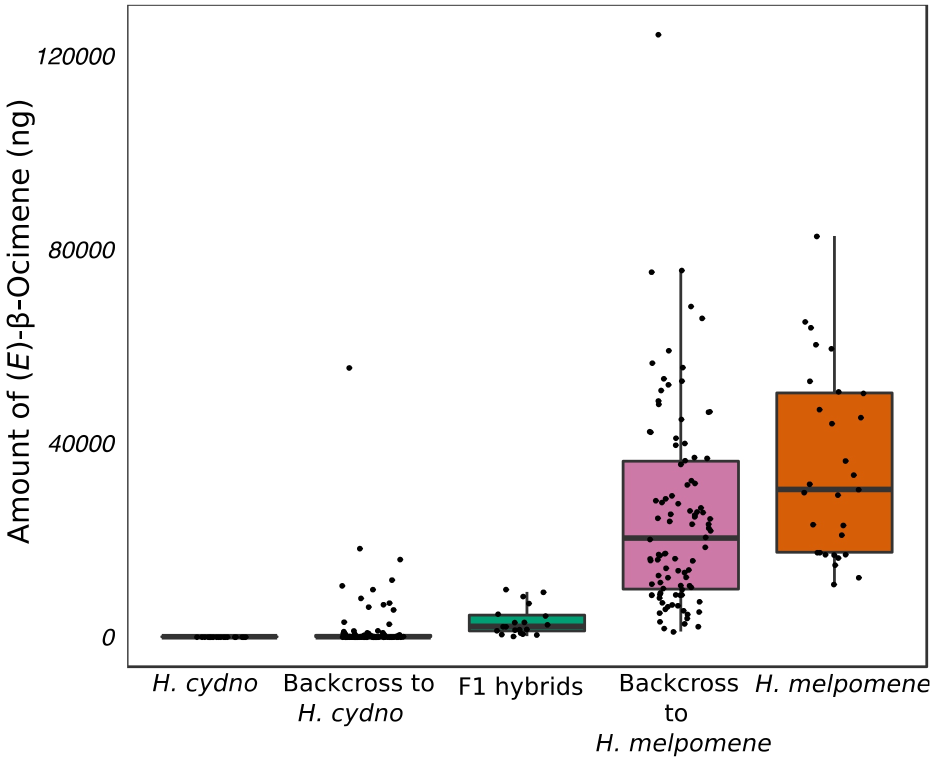

***Figure S2:*** *Effect plot for QTL peak on chromosome 6. Log amount of (*E*)-β-ocimene produced by each genotype at the marker with the highest LOD score. Individuals homozygous for* H. cydno *alleles produce less (*E*)-β-ocimene than heterozygotes with a* H. melpomene *allele.*

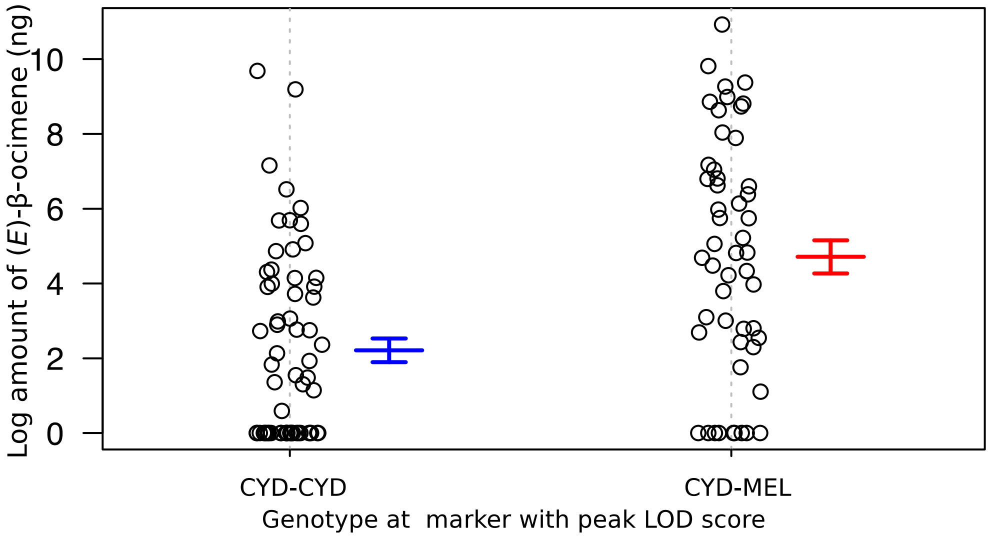

***Figure S3:*** *Log2-expression of the candidate genes in* H. melpomene *and H. cydno abdomens in males and females. females. Both HMEL037106g1 and HMEL037108g1 (highlighted in bold) show greater male-biased expression in* H. melpomene *than* H. cydno. *Full model statistics in Table S4. N=5 for each boxplot. Gene expression is given in the log2 of the normalised counts using TMM (trimmed mean of M values) normalisation.*

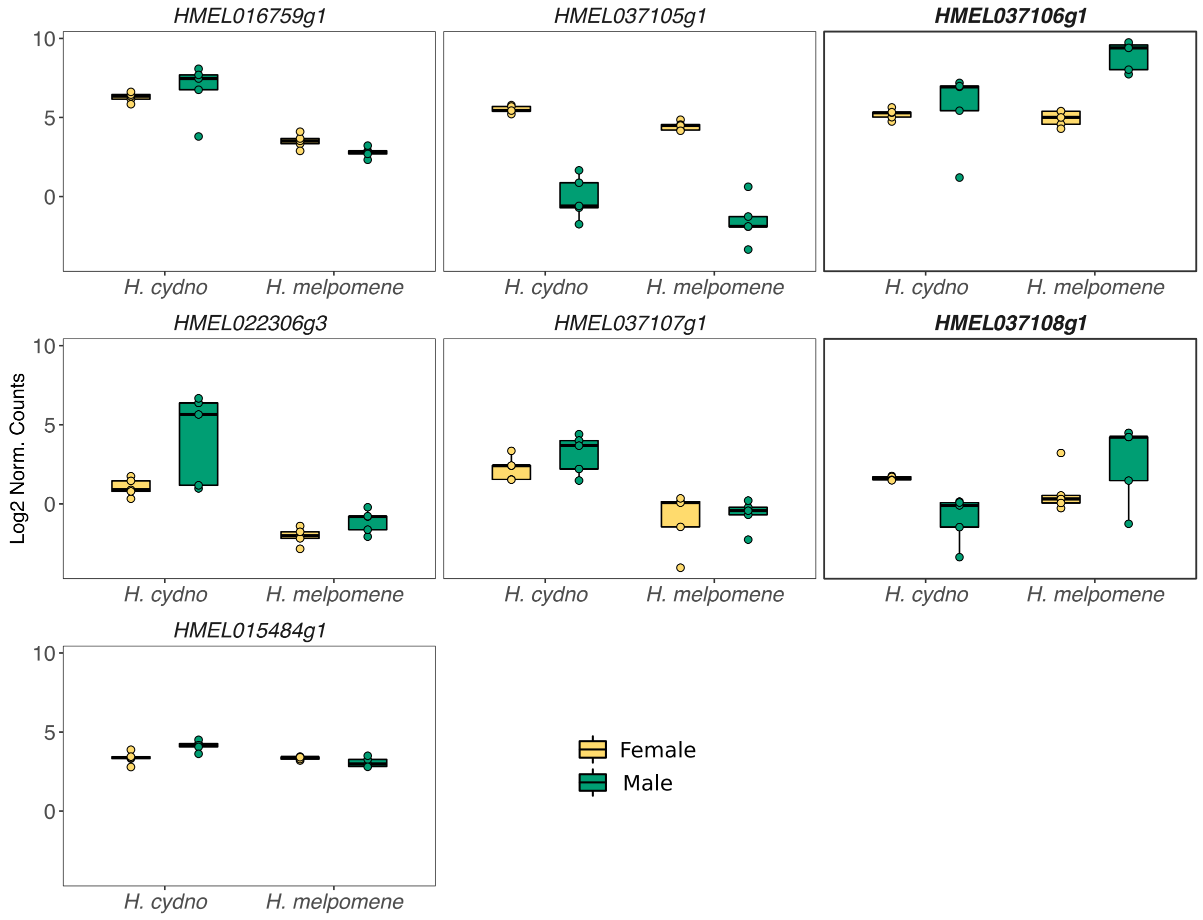

***Figure S4:** Control experiments (protein expression uninduced) for the functional characterisation of TPS activity of A) HMEL037106g1 and B) HMEL037108g1 from* H. melpomene*. Total ion chromatograms of products in the presence of different precursor compounds. (*E*)-β-Ocimene is not produced in any treatments. Linalool and geraniol are produced in small amounts in both, likely due to endogenous bacterial activity. 1, Linalool; 2, Geraniol; *, contaminant from medium; IS, internal standard. Abundance is scaled to the highest peak of all panels per enzyme. Quantification of peaks in Table S6 and S7.*

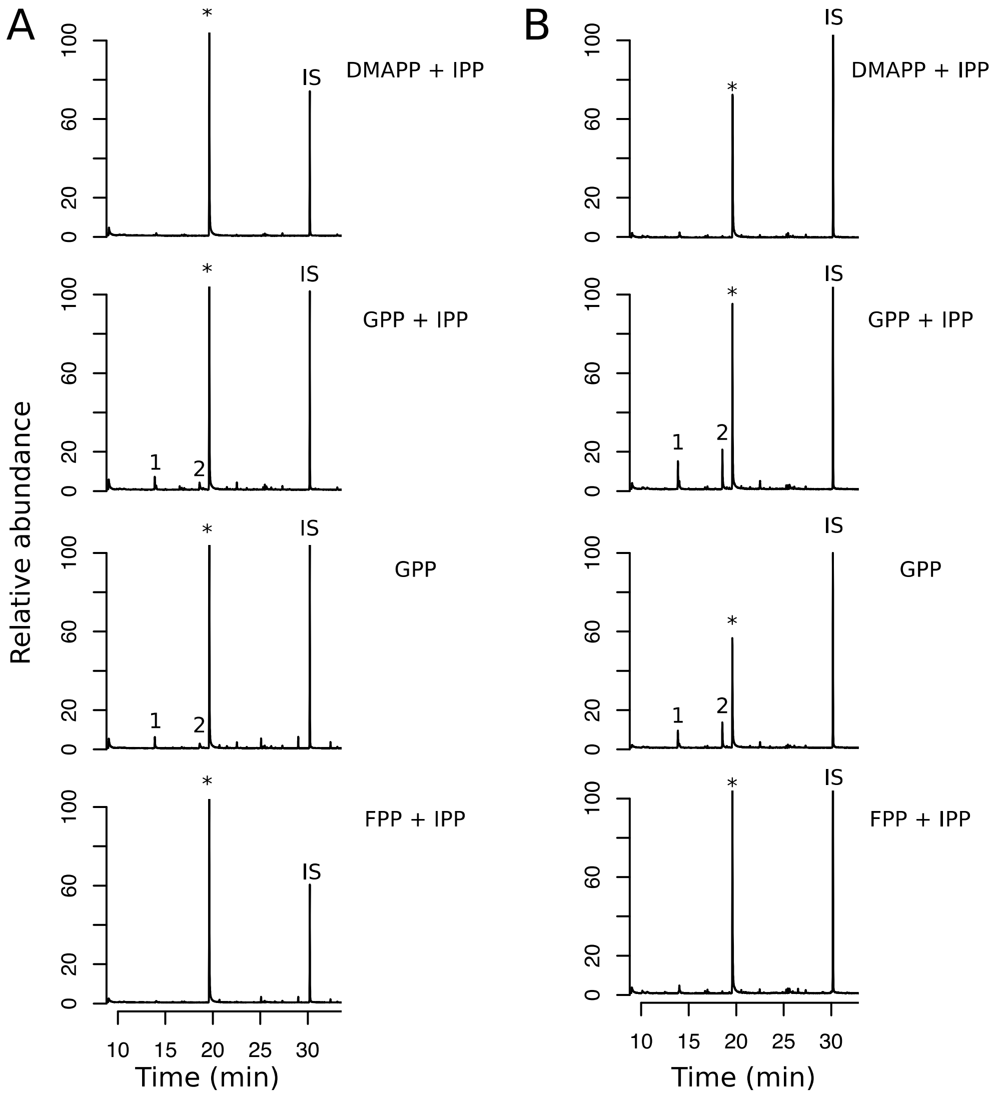

***Figure S5:*** *Linalool is not metabolized into ocimene by HMEL037106g1. A) Total ion chromatograms of enzymatic products in the presence of different linalool stereoisomers. No enzymatic activity is detected. B) Total ion chromatograms of control experiments (protein expression not induced) in the presence of different Linalool stereoisomers. Again, as expected, no enzymatic activity is detected. 1, Linalool; *, contaminants from medium; IS, internal standard. Abundance is scaled to the highest peak of all panels. Quantification of peaks in Table S8.*

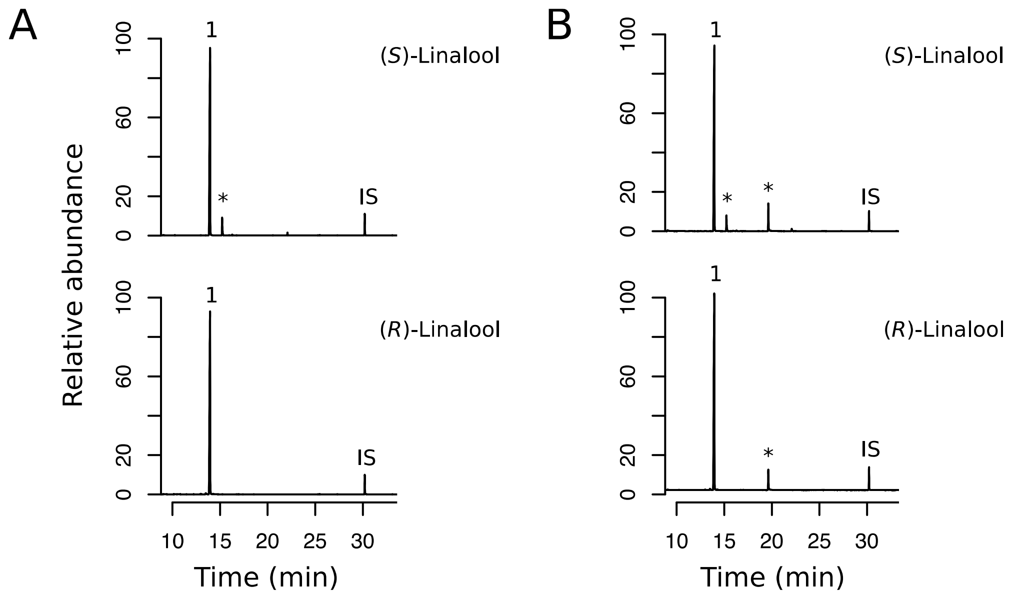

***Figure S6:*** *Chiral analysis of linalool produced by HMEL037106g1 and HMEL037108g1. A) Linalool produced in experiments with HMEL037106g1 is mainly (*S*)-linalool (ratio 97:3,* S:R*), B) linalool produced in experiments with HMEL037108g1 is a racemic mixture (ratio 54:56,* S:R*), C) (*R*)-linalool, D) (*S*)-linalool, E) Racemic linalool mixture.*

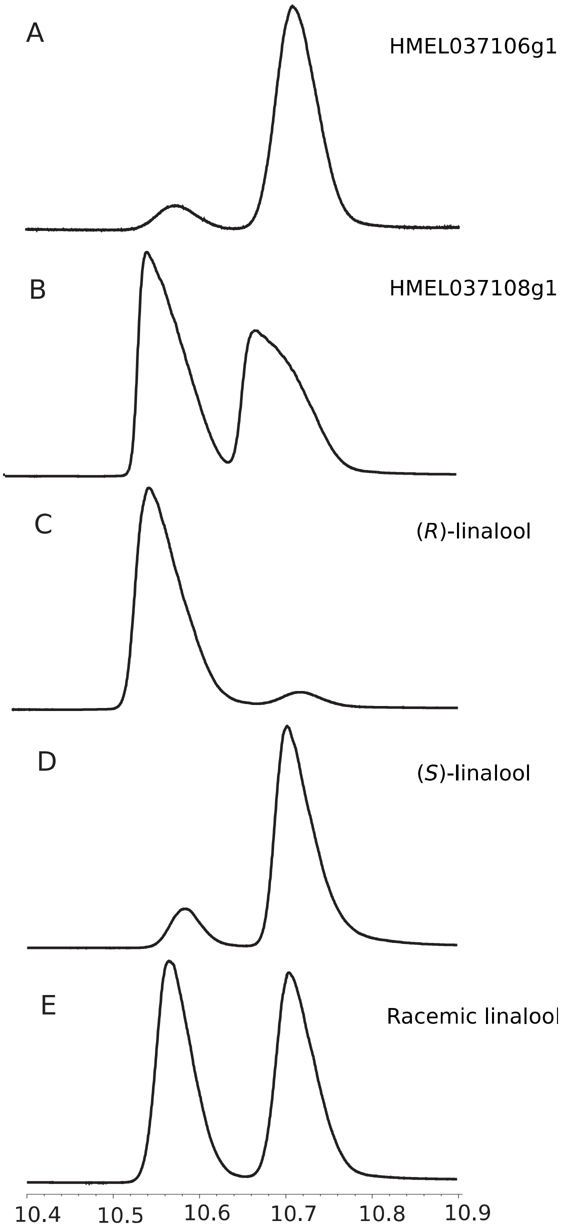

***Figure S7:*** *Functional characterisation of IDS activity of HMEL037106g1 from* H. melpomene. *A) Total ion chromatograms of enzymatic products in the presence of different precursor compounds, following treatment by alkaline phosphatase. GPP is dephosphorylated to produce geraniol, and FPP to produce farnesol, demonstrating that the main function of HMEL037106g1 is not as an IDS. B) Total ion chromatograms of control experiments (protein expression not induced) in the presence of different precursor compounds, following treatment by alkaline phosphatase. As expected, GPP is dephosphorylated to geraniol, and FPP to farnesol. 1, (*E*)-*β*-Ocimene; 2, Linalool; 3, Geraniol; 4, Farnesol; *, contaminant from medium; IS, internal standard. Abundance is scaled to the highest peak of all panels. Quantification of peaks in Table S10.*

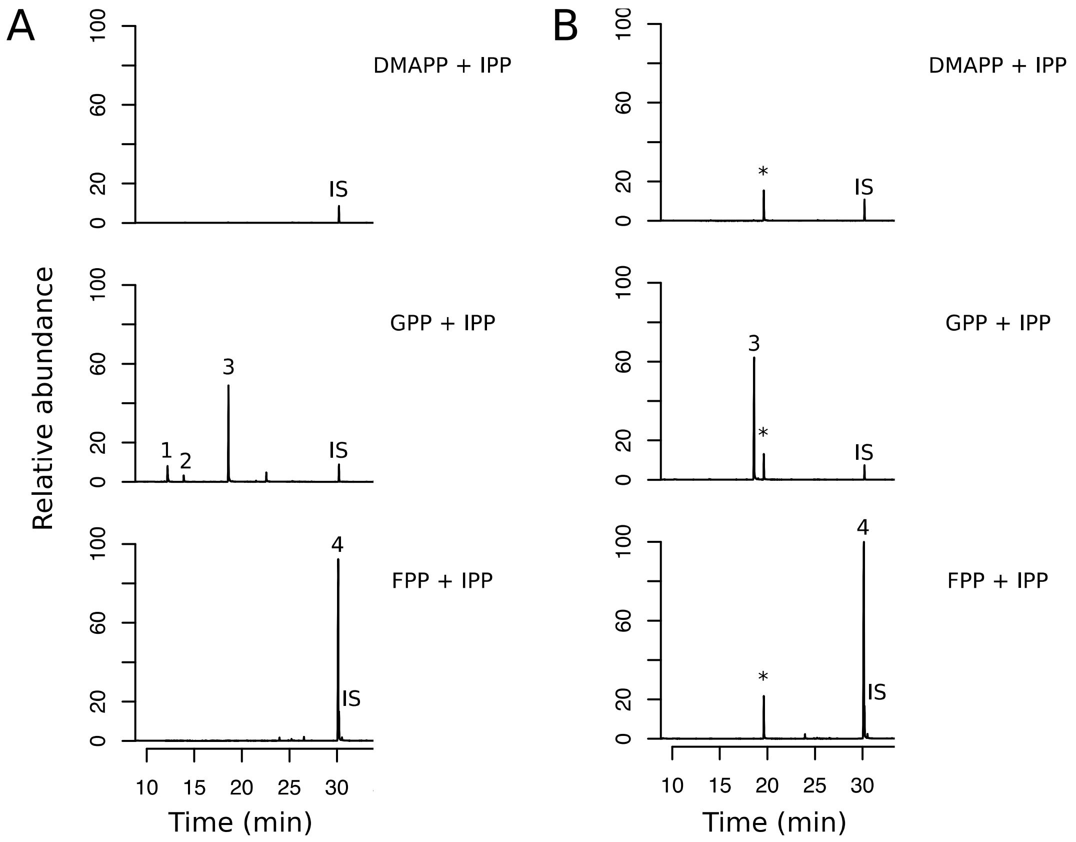

***Figure S8:*** *Functional characterisation of IDS activity of HMEL037108g1 from* H. melpomene. *A) Total ion chromatograms of enzymatic products in the presence of different precursor compounds, following treatment by alkaline phosphatase. As in Figure S6, GPP is converted to linalool and FPP to nerolidol, with remaining GPP dephosphorylated to geraniol, and FPP to farnesol. HMEL037108g1 is acting as a mono- and sesquitepene synthase, not an IDS. B) Total ion chromatograms of control experiments (protein expression not induced) in the presence of different precursor compounds, following treatment by alkaline phosphatase. GPP is dephosphorylated to geraniol and FPP to farnesol. 1, Linalool; 2, Geraniol; 3, Nerolidol; 4, Farnesol; *, contaminant from medium; IS, internal standard. Abundance is scaled to the highest peak of all panels. Quantification of peaks in Table S11.*

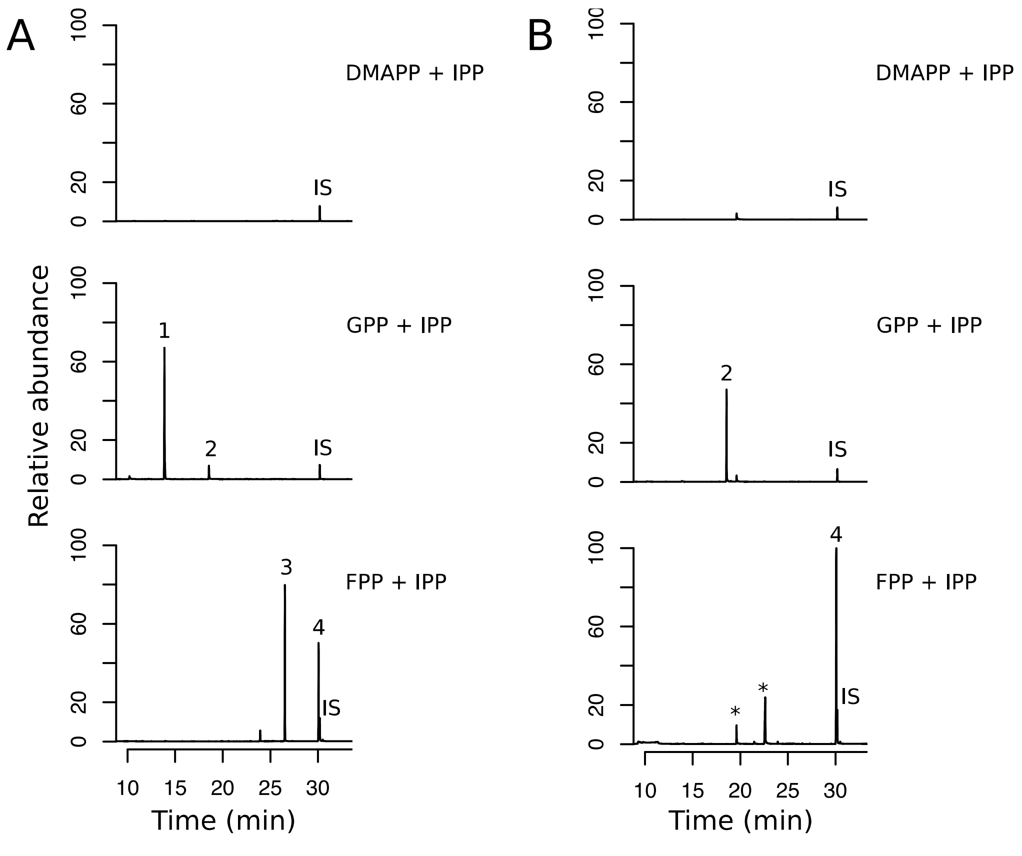

***Figure S9:*** *Unrooted phylogenetic tree showing the relationships between protein sequences of GGPPSs in Lepidoptera. The tree was obtained from FastTree, a tool for creating approximately-maximum-likelihood trees, using the JTT (Jones-Taylor-Thornton) model of amino acid evolution. Local support values are illustrated.*

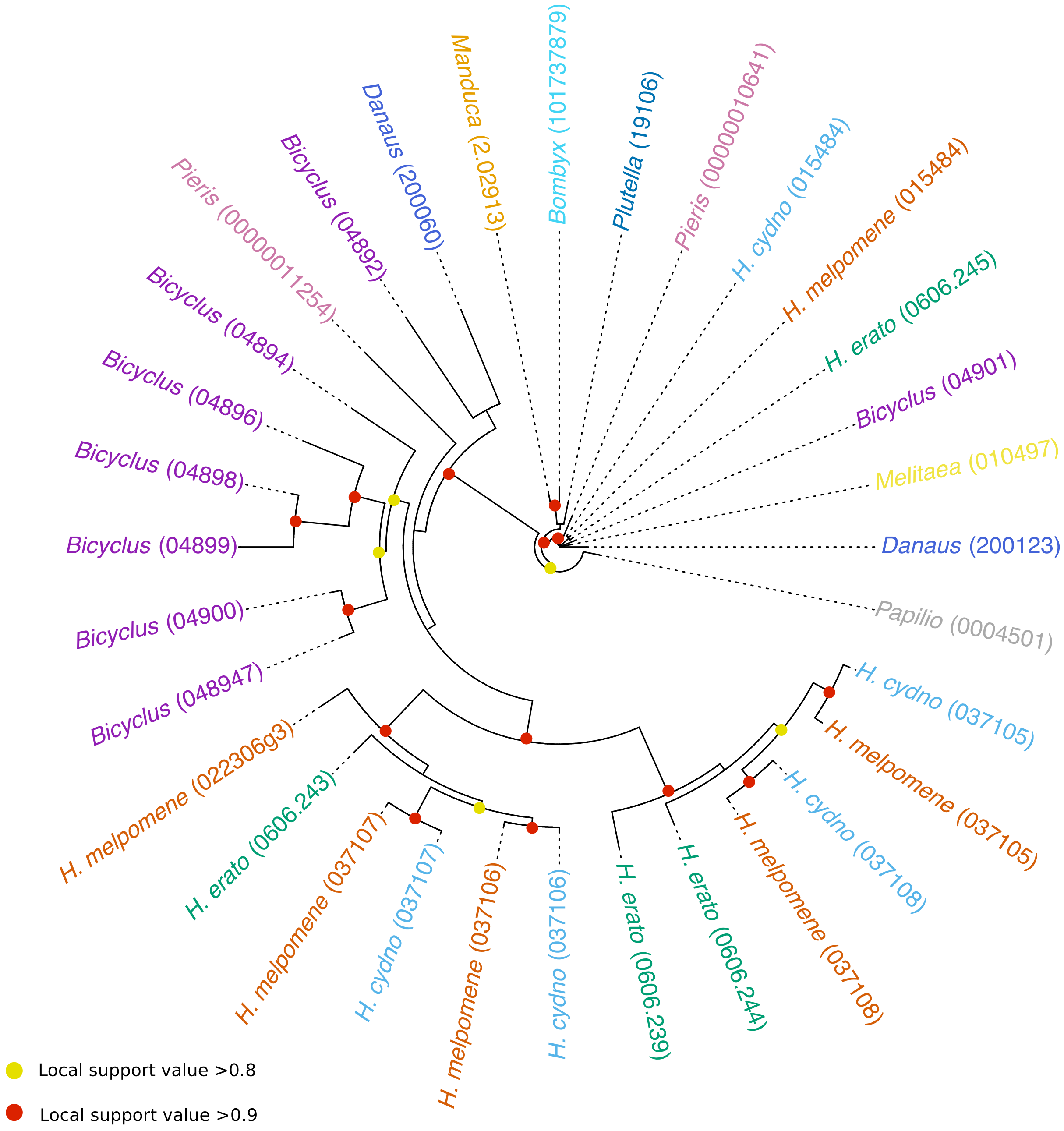

***Figure S10:*** *Phylogram of genes annotated as GGPPSs in* Heliconius melpomene*,* H. cydno*, and* H. erato*, including* HMEL037106g1 *and* HMEL037108g1 (*) *which act as TPSs. The phylogeny was constructed in FastTree using the Jukes-Cantor model of nucleotide evolution. Well-supported branches are illustrated. The* H. erato *gene* Herato0606.245 (GGPPS, shows high similarity to the GGPPS of the moth *Choristoneura fumiferana*) *was used to root the tree.*

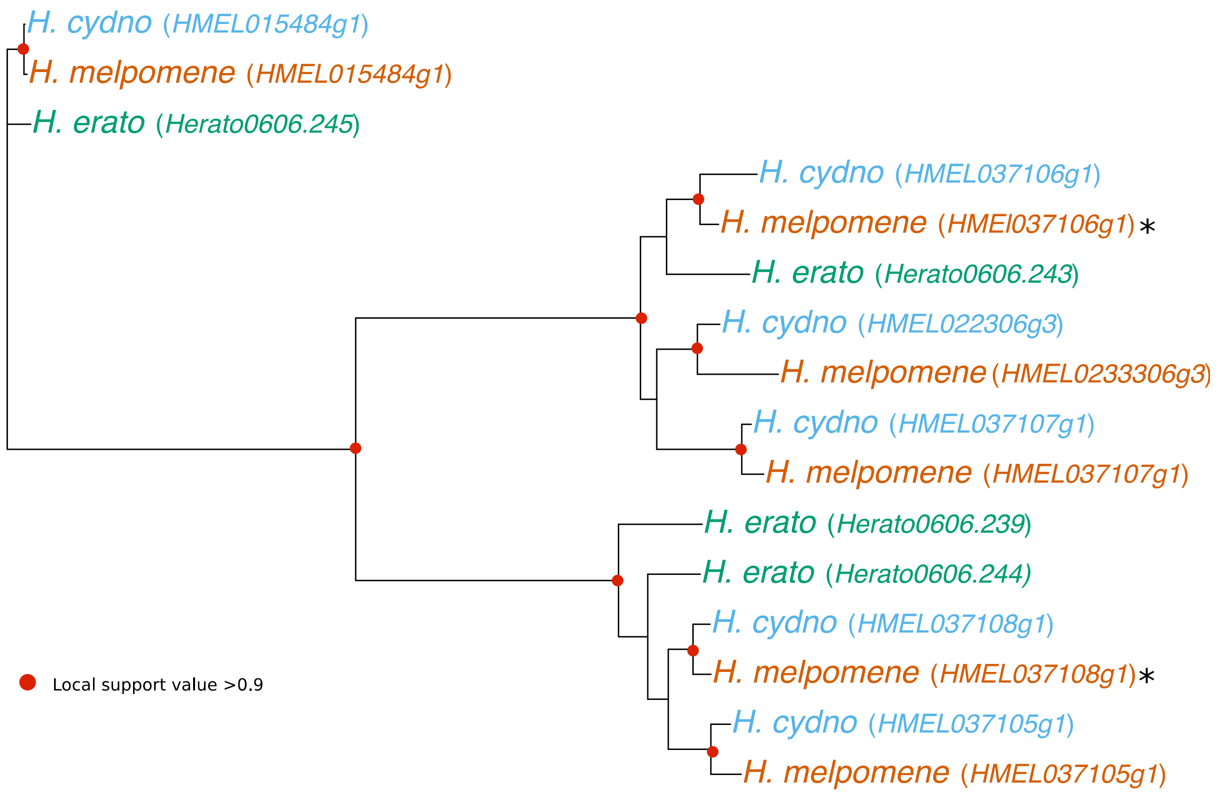

***Figure S11:*** *Amino acid alignment between* H. melpomene *GGPPS and TPSs, and other insect TPSs. The two aspartate-rich motifs are labelled FARM and SARM. Stars show residues identified as conserved between insect TPSs (17). Hmel,* H. melpomene*; Mhistr,* Murgantia histrionica*; Ip*, Ips pini*; Pstri,* Phyllotreta striolata*; Hmel_TPS_1, HMEL037106g1; Hmel_TPS_2, HMEL037108g1.*

```
Hmel_GGPPS    1 -------------------------------------------------------MSLV-
Hmel_TPS_2    1 -------------------------------------------------------MDVQK
Hmel_TPS_1    1 ---------------------------MSE---------------------TEVHVVKVN
Mhistr_TPS    1 ------------MVSIAAKSLPKLSGAVFG-------------------------QFSRR
Ip_IDS_TPS    1 MFKLAQR---------LPKSVSSLGSQLSKNAPNQLAAATTSQLINTPGIRHKSRSSAVP
Pstri_TPS3    1 --MLYVLKNYNLNYSIVSKVPLHFRSLC----------------------------SLLQ
Pstri_TPS4    1 MFAICKVVNYYSSCRIIPKVSGNFGTLLQRSF---------------------NRAFSCE
Pstri_TPS1    1 MFLLPRLKNFTRSNSPARKLFSPK----SN-------------------------SFS--
Pstri_TPS2    1 ------------------------------------------------------------

Hmel_GGPPS    5 KSKDGDKNQD------------------EKLLMP--FTYIQQVPGKQIRSKLTLAFNYWL
Hmel_TPS_2    6 ISENSDLYME------------------NELLLP--YNHVLQVSGKQMRMKIVKALNHWL
Hmel_TPS_1   13 GKENDDLFLE------------------KEILAP--FSHICQVKGKQLRIKIMRAFNHWL
Mhistr_TPS   24 KQLIQRHWL--------DTRTDQYYDVLRRIVVPECKNIASDVPEYPERIEKLLYYTNPA
Ip_IDS_TPS   52 SSLSKSMYDHNEEMKAAMKYMDEIYPEVMGQIEK--VPQYEEIKPILVRLREAIDYTVPY
Pstri_TPS3   31 KK-NNRPLVD--I-SVEEGPLRSVYPAIREEIIEEHLVLK-GNSELRDRCEKLLDYNANV
Pstri_TPS4   40 IE-ANEPFVD--L-FSEEEHLKSMLPAVKEEIIEEHLVLHKDNKEIRNRCEKLIDYNINT
Pstri_TPS1   30 ----STPHDD-GFFKHEMDELKTYYPLMVQDLTDA-ISQYKQFPGLLERFPVLMDYTVTH
Pstri_TPS2    1 ------MYTQ-DFFLDEYNEIMAYYPKVIEQVRN--WTFIKEYQGLPDRYTNYMNYNVDV
                                                                 FARM
Hmel_GGPPS   45 -----------------RV------SDD---KLRAIGEIVQMLHNSSLLLDDIQDNSILR
Hmel_TPS_2   46 -----------------KV------PEG---DMENIVNLIHMIHAASLLLDDIQDDSKLR
Hmel_TPS_1   53 -----------------QA------SEV---DVMKALGIVNSLHVASLLIDDVQDDSTLR
Mhistr_TPS   76 F---SDAWNFTTELIYRTVADESHQTEENITKMYLIRATMDLLFTMSAVLDDISDRSEFR
Ip_IDS_TPS  110 GKR-F--KGVHIVSHFKLLADPKFITPENVKLSGVLGWCAEIIQAYFCMLDDIMDDSDTR
Pstri_TPS3   86 ETP-FLSASLIFLHTYKLLEKPSLLNHENLKKAYILAWCHKLIHSSININDDIIDRSNIR
Pstri_TPS4   96 ESK-FLTFPLIFLRTYKLLEKPALLNDENLKKACILAWCHRLIHASVIISDDIVDDSEMR
Pstri_TPS1   84 DDPYFLSSAVLPLYFYKAVEESDKLTEENIKRACLMSWAYRTLETSQIIVDDILDKSEVR
Pstri_TPS2   52 KDTKRMATALFTLYSYKHLEQPEKQTDENLRKAIAMAWAFRMAEASQLTLDDVLDNSLTR

Hmel_GGPPS   79 RGIPVAHSIYGIAST--INAANY---VIIIALEKTLKLGHPLA---TTVYTEQLLELHRG
Hmel_TPS_2   80 RGLPAAHNVYGVPLT--LNASCH---AIFLVLIKSYDIN-PKV---SKIMVESFLWGLRG
Hmel_TPS_1   87 RGMPAAHCVYGVPLT--VNTSLH---AMFLVLEEAFAVD-PKA---AKLLVEDFLEMCRG
Mhistr_TPS  133 KGKKGWHMICQGGESTALYDGTQMGLFPLYLLKQY-FKNDPGYSRLLETVVMTYIKLTIG
Ip_IDS_TPS  167 RGKPTWYKLPGIGLN-AVTDVCLMEMFTFELLKRY-FPKHPSYADIHEILRNLLFLTHMG
Pstri_TPS3  145 YNKTTWYQLPDVGKDAAIVDAAFLLNGAIFLLQNH-LRCLPHQYIMQKHFLRAHAIMNLS
Pstri_TPS4  155 YNKTTWCKLPDVGKEDAITDAAFLLTGAIFLLQNH-LRHHPHNFILQKHFLRGYAFINVS
Pstri_TPS1  144 YNKPAWYKKDGVSMELTILDSHYLATGAYMVLTKR-LAGHPCCLDILDLYAEEMFVMIIA
Pstri_TPS2  112 YMKPAWHKLEGTNN--AVLDAFFVENAAYLILQEE-MRDHPQFLNIVKLLKEYYLMLVVG

Hmel_GGPPS  131 QGLEIYWR------DNFQCPTEDEYKEMTMKKT--GGLFM-----LAIRLMQLFSDNKS-
Hmel_TPS_2  131 QGMDLHWR------ENFVCPTVEQYMKMVELKT--GYMFS-----GAYEVMQLYSDNKT-
Hmel_TPS_1  138 QGIDIYWR------DHLICPTEGQYYKMLEQKT--GHFFL-----MGVRMMQLFSCNKT-
Mhistr_TPS  192 QTIDV------LGQFKKSPSM-AEYKRINYYKA--GQFVA-AGSELAVIHAGITSQDLID
Ip_IDS_TPS  225 QGYDFTFIDPVTRKINFNDFTEENYTKLCRYKIIFS--TFHNTLELTSAMANVYDPKKIK
Pstri_TPS3  204 NIPE-------LKKIKIN---E-----GDKHQLDIKFYSYN-IISTAMFLANVTDGYLHE
Pstri_TPS4  214 YMMD-------NRKHHIN---E-----LEKYQVHTKLYFSN-LFSTAMYLANVENDYWQK
Pstri_TPS1  203 QYMD-------IKKLDLKDFQK-----LVRHRFDKALYVFNGSARSGLYLANVRDRETHD
Pstri_TPS2  169 QYLD-------MRSIPFEKSFE-----LLKYRNIKGYYITNMPIRGSMYLANIDNPDYHA
                                 SARM
Hmel_GGPPS  177 DFTKLSSLLGLYFQIRDDYCNLCLREY------------SENKSYCEDLTEGKFSFPIIH
Hmel_TPS_2  177 DYSKIISIMGRLFQIRDDYCNLKHREALEEWPGEDDIEVIRDHDFCEDITEGKFSFPVIH
Hmel_TPS_1  184 DYSELVLLMGRYFQIRDDYCNLSQQEALEEWPGAEDLQVCKNDSFCEDITEGKISLPIIH
Mhistr_TPS  242 KTVEIFTIAGQIIQTWDDFNDYYSSSEQN-------GK------LSCDFMNAGTTWVSAK
Ip_IDS_TPS  283 QLDPVLMRIGMMHQSQNDFKDLYRDQGEV-------LKQAEKSVLGTDIKTGQLTWFAQK
Pstri_TPS3  248 VCEEICNDLSRFIRIEDDVVDLYDSEGNI-------RKT-----SCTDISLGRPSWLTME
Pstri_TPS4  258 VSDEICNDISQYLKVEDDVIDLYDSKGKI-------R-------KCTDISLGRPSWFLME
Pstri_TPS1  251 CMKKFSVPMSRFFQVQNDFSGVFEEESKF-------Q-N-----SCPDIVNGRNSWLVTT
Pstri_TPS2  217 KVEEILRISGEWIIIQNDYQEVFLPTSEN-------K-K-----DRRDIQQGTNTWCLAK
```

```
Hmel_GGPPS  225 AIQNQKGDN-------------QVLHILRQRTRDVEVKRYCISLLEKFGSFQYTRDCLQE
Hmel_TPS_2  237 ALSTPEGK--------------PILNILKQRTRDVQLKKYCVSLMEKIGSLQHTCDVLDK
Hmel_TPS_1  244 ALQTKKAG--------------IVMNILRQKTRDMYLKKYCVSTLEEIGSLQYTRNVLEK
Mhistr_TPS  289 AMEVFTPSQAVKFMECYGSDDQSKMKTVQELYDEIDMPKLYTEYVLEN---------Y--
Ip_IDS_TPS  336 ALSICNDRQRKIIMDNYGKEDNKNSEAVREVYEELDLKGKFMEFEEES---------F--
Pstri_TPS3  296 AYKKGSAAQKKILEENFGKNNEESTEKIYSIFEDLQLLDVYKKLSDEF---------Y--
Pstri_TPS4  304 AYKRANAGQRKILEENFHKNNEESVEKLYSIFQELELLEVYRKFTDNF---------Y--
Pstri_TPS1  298 ALKMANPAQRKVIEENYGNGDAESARKVMQVYEDLKLKDVHDRRTEEF---------L--
Pstri_TPS2  264 ALELASESQMKVLKENYGKNDDESAMKIEEIYRDLKLDEIYLKIEEEY---------F--

Hmel_GGPPS  272 LDNEARAEVQRLGGNPHLEDLLDELLSWREDKKSAVNEE------------
Hmel_TPS_2  283 LDQEAREEVARLGGNPEMIAVLDELLSWKTN--------------------
Hmel_TPS_1  290 LDLEIRAEVARLGGNPLIDEVLHSLLSWKDN--------------------
Mhistr_TPS  338 --NRCETLIKEL-PHDRLREACSSYMEWLVVRETPDEDSEHKVALCLNISG
Ip_IDS_TPS  385 --EWLKKEIPKI-NNGIPHKVFQDYTYGVFKRRPE----------------
Pstri_TPS3  345 --EQAI---YKI-QKKLPKSKMQDALLDLLTLIVNHKCM------------
Pstri_TPS4  353 --AQEI---SKI-REKIPKSIMQDILINLVNLAVNHKLRY-----------
Pstri_TPS1  347 --GEMREIVENF-PERIPKQPFHDIVRQLALNKLYS---------------
Pstri_TPS2  313 --EKVNRRLDIL-PNILPKSFFWNMMHIIKNEYMNG---------------
```

***Table S1:*** Drosophila melanogaster *query protein sequences downloaded from FlyBase and searched (blastp) against all annotated proteins in the genome of* Heliconius melpomene *(v2.5) on LepBase to identify homologs of enzymes involved in the mevalonate and putative terpene synthesis pathways. The candidate orthologs identified in* H. melpomene *were then searched (blastp) against annotated proteins in the* D. melanogaster *genome on FlyBase. Reciprocal best blasts are highlighted in bold. We included other hits with an e-value smaller than* $1e^{-80}$.

| Gene (symbol) | *D. melanogaster* | *H. melpomene* |
|---|---|---|
| Acetoacetyl-CoA thiolase (ACAT2) | CG9149 | HMEL032609g1<br>HMEL014614g2<br>HMEL017484g1 |
| Hydroxymethylglutaryl-CoA synthase (HMGCS) | CG4311 | **HMEL005451g1** |
| Hydroxymethylglutaryl-CoA reductase (HMGCR) | CG10367 | **HMEL016133g1** |
| Mevalonate kinase (MVK) | CG33671 | **HMEL013262g2** |
| Phosphomevalonate kinase (PMVK) | CG10268 | **HMEL007429g2/3*** |
| Diphosphomevalonate decarboxylase (MVD) | CG8239 | **HMEL004012g1** |
| Isopentenyl-diphosphate isomerase (IDI) | CG5919 | **HMEL005103g1** |
| Farnesyl pyrophosphate synthase (FPPS) | CG12389 | **HMEL017961g1**<br>HMEL017961g2 |

| | | |
|---|---|---|
| Geranylgeranyl pyrophosphate synthase (GGPPS) | CG8593 | **HMEL015484g1**<br>HMEL037105g1<br>HMEL037106g1<br>HMEL022306g3<br>HMEL037107g1<br>HMEL037108g1 |
| Decaprenyl pyrophosphate synthase subunit 1 (PDSS1) | CG31005 | **HMEL016759g1** |
| Decaprenyl pyrophosphate synthase subunit 2 (PDSS2) | CG10585 | **HMEL031784g1**<br>HMEL011234g1<br>HMEL008172g1<br>HMEL008173g1 |

*The first two exons of *HMEL007429g2* and the last exon of *HMEL007429g3* are expressed as a single transcript: (ATGGCACCAAAGATTGTATTATTGTTCAGTGGGAAAAGAAAATGTGGTAAAGACTTCGTGACTGATCATCTTAAGACATTGTTAAGTGACCAATGTGAAATTATAAAAATTTCACAACCCATCAAAAGTCATTGGGCAAAGGAAAAGAAATTAAATTTAAATGATCTCTTAAGTGATGGTGAATATAAAGAGAACTACCGCCTAGAAATGATAAAATGGAGTGAGGAAATGAGACAAAAAGATTATGGTTGTTTTTGTAGAGCTGCATGTGAAAATGCTACAGAGAAACCTGTATGGATTGTCAGTGATATAAGACGGAAAACAGATTTGCAGTGGTTTAAAGAAACCTATGGTGATCTTATTAAAACAATTCGACTAACAGCAGATGACAATACTAGGACTGAAAGAGGTTTCCAATTTAAGAGTGGAGTTGATAATGCAGCCTCAGAATGTGATTTAGACGATTATACAGAATGGGATCTCATTATTGAAAACAGCAAAGATAAAACTGTTGAGGATCTTACTAAAAATATTATACTTCTATTAGAATCTTTGAATATTTTACACAAACTTAGGTAG)

***Table S2:*** *Transcript sequences for* H. cydno *genes from QTL region, as well as names used in* H. cydno *annotation. (Attached excel file).*

**Table S3:** *Linear model statistics for differential gene expression analysis in* H. melpomene *heads and abdomens of both sexes. The model includes two fixed terms, tissue and sex, their interaction, and a random term, individual (expression ~ sex + tissue + sex*tissue + (1|individual). The Log FC column gives the log2 Fold Change between the groups being compared, while the Ave. Expr. column gives the mean log2-exprtession across all samples. Column t is the moderated t-statistic and B is the B-statistic, the log odds that the gene is differentially expressed. The Adj. p-value column gives p-values (bold are significant) corrected for multiple testing using the Benjamini and Hochberg's method to control the false discovery rate across all tested genes (17,902).*

| Gene | Term | Log FC | Ave. Expr. | t | p-value | Adj. p-value | B |
|---|---|---|---|---|---|---|---|
| *HMEL016759g1* | sex*tissue | 0.6677 | 4.3438 | 1.3668 | 0.1863 | 0.4143 | -6.1253 |
| | sex | -0.7177 | 4.3438 | -1.5627 | 0.1333 | 0.3131 | -5.9173 |
| | tissue | 2.0102 | 4.3438 | 6.1544 | 4.41E-06 | **3.14E-05** | 3.7935 |
| *HMEL037105g1* | sex*tissue | 2.9238 | -0.6946 | 2.4561 | 0.0230 | 0.0962 | -3.7249 |
| | sex | -6.1677 | -0.6946 | -7.2360 | 4.27E-07 | **5.36E-06** | 6.5817 |
| | tissue | -5.3853 | -0.6946 | -7.5482 | 2.24E-07 | **2.42E-06** | 7.1670 |
| *HMEL037106g1* | sex*tissue | -4.8560 | 3.0924 | -4.3466 | 0.0003 | **0.0028** | 0.2480 |
| | sex | 3.9182 | 3.0924 | 7.6856 | 1.69E-07 | **2.26E-06** | 7.1353 |
| | tissue | -5.0244 | 3.0924 | -6.6000 | 1.65E-06 | **1.36E-05** | 5.1304 |
| *HMEL022306g3* | sex*tissue | -1.7761 | -2.6439 | -2.0475 | 0.0535 | 0.1795 | -4.3872 |
| | sex | 1.0489 | -2.6439 | 1.4821 | 0.1534 | 0.3461 | -5.4490 |
| | tissue | -0.8095 | -2.6439 | -1.3163 | 0.2024 | 0.2995 | -5.7741 |
| *HMEL037107g1* | sex*tissue | -1.5469 | -1.2854 | -1.4056 | 0.1747 | 0.3976 | -5.4444 |
| | sex | 0.2268 | -1.2854 | 0.2483 | 0.8063 | 0.8941 | -6.5585 |
| | tissue | 0.0553 | -1.2854 | 0.0731 | 0.9424 | 0.9609 | -6.7482 |
| *HMEL037108g1* | sex*tissue | -1.9477 | 1.1227 | -1.7246 | 0.0995 | 0.2750 | -5.2255 |
| | sex | 2.0789 | 1.1227 | 2.3922 | 0.0263 | 0.0953 | -4.2979 |
| | tissue | -0.2850 | 1.1227 | -0.3439 | 0.7344 | 0.8065 | -6.8774 |
| *HMEL015484g1* | sex*tissue | 0.4644 | 4.0455 | 1.1758 | 0.2530 | 0.5043 | -6.3477 |
| | sex | -0.2477 | 4.0455 | -0.6846 | 0.5012 | 0.6873 | -6.8918 |
| | tissue | 1.3734 | 4.0455 | 5.0788 | 0.0001 | **0.0003** | 1.3380 |

**Table S4:** *Linear model statistics for differential gene expression analysis in* H. melpomene *and* H. cydno *abdomens of both sexes. The model includes two fixed terms, species and sex, and their interaction (expression ~ sex + species + species*tissue). The Log FC column gives the log2 Fold Change between the groups being compared, while the Ave. Expr. column gives the mean log2-exprtession across all samples. Column t is the moderated t-statistic and B is the B-statistic, the log odds that the gene is differentially expressed. The Adj. p-value column gives p-values (bold are significant) corrected for multiple testing using the Benjamini and Hochberg's method to control the false discovery rate across all tested genes (11,571).*

| Gene | Term | Log FC | Ave. Expr. | t | p-value | Adj. p-value | B |
|---|---|---|---|---|---|---|---|
| *HMEL016759g1* | species*sex | -0.3223 | 5.4126 | -1.4740 | 0.1558 | 0.4349 | -6.6939 |
| | species | 1.7233 | 5.4126 | 7.8810 | 1.32E-07 | **4.59E-07** | 6.6022 |
| | sex | 0.0672 | 5.4126 | 0.3071 | 0.7619 | 0.8445 | -8.1198 |
| *HMEL037105g1* | species*sex | -0.1478 | 2.5652 | -0.7547 | 0.4591 | 0.7264 | -7.0952 |
| | species | 0.6662 | 2.5652 | 3.4010 | 0.0028 | **0.0061** | -3.1192 |
| | sex | 2.9925 | 2.5652 | 15.2781 | 1.33E-12 | **3.14E-11** | 18.9110 |
| *HMEL037106g1* | species*sex | 0.8480 | 6.7288 | 3.1511 | 0.0050 | **0.0446** | -3.7029 |
| | species | -0.7377 | 6.7288 | -2.7414 | 0.0125 | **0.0236** | -5.1106 |
| | sex | -1.0794 | 6.7288 | -4.0113 | 0.0007 | **0.0027** | -2.1200 |
| *HMEL022306g3* | species*sex | -0.6182 | 0.9778 | -1.3147 | 0.2033 | 0.4950 | -6.4067 |
| | species | 2.2782 | 0.9778 | 4.8446 | 0.0001 | **0.0003** | 0.3653 |
| | sex | -1.1116 | 0.9778 | -2.3639 | 0.0282 | 0.0699 | -5.0829 |
| *HMEL037107g1* | species*sex | -0.2153 | 1.4275 | -0.7773 | 0.4459 | 0.7173 | -7.0103 |
| | species | 1.8967 | 1.4275 | 6.8481 | 1.09E-06 | **3.53E-06** | 4.8657 |
| | sex | -0.3254 | 1.4275 | -1.1747 | 0.2537 | 0.3944 | -7.0160 |
| *HMEL037108g1* | species*sex | 1.1927 | 1.5270 | 3.4427 | 0.0025 | **0.0257** | -2.4568 |
| | species | -0.8135 | 1.5270 | -2.3483 | 0.0291 | **0.0503** | -5.2982 |
| | sex | 0.1373 | 1.5270 | 0.3963 | 0.6960 | 0.7963 | -7.6400 |
| *HMEL015484g1* | species*sex | -0.2868 | 4.0561 | -3.8325 | 0.0010 | **0.0123** | -1.9266 |
| | species | 0.2741 | 4.0561 | 3.6640 | 0.0015 | **0.0035** | -2.8338 |
| | sex | -0.1316 | 4.0561 | -1.7586 | 0.0937 | 0.1854 | -6.5949 |

***Table S5:*** *Primer sequences. CACC was added to the 5' end of the forward primer so that it was compatible with the plasmid vector.*

| Gene | Primer sequence | Use |
|---|---|---|
| *HMEL037106g1* | Forward: CACCATGTCAGAAACAGAAGTCC Reverse: TTAATTATCCTTCCAACTTAAAAGCGA | Amplification of transcript from cDNA library |
| *HMEL037108g1* | Forward: CACCATGGACGTTCAGAAAATAAGC Reverse: TTAATTCGTTTTCCAAGAAAGAAGTTC | Amplification of transcript from cDNA library |
| *HMEL037106g1* | Forward: GTTAATTCGTTACACGTAGC Reverse: TAATCTGGAAATAGCGACC | Sequencing PCR products |
| *HMEL037108g1* | Forward: CTAACTCTTAACGCCTCG Reverse: TTATATCCTCGCAGAAATCG | Sequencing PCR products |
| Both | Forward: AATACGACTCACTATAGGGG Reverse: GGTTAGGGATAGGCTTACC | Sequencing insert in plasmid |

***Table S6:*** *Quantification of experiments characterising TPS activity of HMEL037106g1 (Fig. 5A, Fig. S4). HMEL037106g1 is a monoterpene synthase, catalysing the formation of (*E*)-β-ocimene from GPP. Residual IDS activity is shown by the production of (*E*)-β-ocimene, linalool, and nerolidol from DMAPP and IPP. Mean amounts (ng) ± standard deviation for each compound across 3 replicates are shown. (control) indicates experiments where protein expression was not induced. N=3 for each treatment.*

| | (*E*)-β-Ocimene | (*Z*)-β-Ocimene | Linalool | Geraniol | Nerolidol |
|---|---|---|---|---|---|
| DMAPP + IPP | 7.8±1.4 | 0±0 | 3.4±0.4 | 0±0 | 4.4±1.8 |
| DMAPP + IPP (control) | 0±0 | 0±0 | 0±0 | 0±0 | 0±0 |
| GPP + IPP | 334.7±32.7 | 10.7±1.5 | 84.2±7.7 | 36±5.2 | 0±0 |
| GPP + IPP (control) | 0±0 | 0±0 | 17.2±2.6 | 17.4±4.6 | 0±0 |
| GPP | 356.5±115.3 | 12.0±4.4 | 108.4±37.3 | 66.6±23 | 0±0 |
| GPP (control) | 0±0 | 0±0 | 10.4±9 | 18.5±6 | 0±0 |
| FPP + IPP | 0±0 | 0±0 | 0±0 | 0±0 | 12.3±0.8 |
| FPP + IPP (control) | 0±0 | 0±0 | 0±0 | 0±0 | 2.1±0.3 |

***Table S7:*** *Quantification of experiments characterising TPS activity of HMEL037108g1 (Fig. 5A, Fig. S4). HMEL037108g1 acts as a mono- and sesquiterpene synthase, producing linalool from GPP and nerolidol from FPP. Small amounts of linalool and nerolidol detected in DMAPP and IPP treatment, and of nerolidol in the GPP treatment, demonstrate residual IDS activity. Mean amounts (ng) ± standard deviation for each compound across 3 replicates are shown. N=3 for each treatment.*

| | (*E*)-β-Ocimene | Linalool | Geraniol | Nerolidol | Farnesol |
|---|---|---|---|---|---|
| DMAPP + IPP | 0±0 | 8.5±0.5 | 0±0 | 10.2±0.5 | 0±0 |
| DMAPP + IPP (control) | 0±0 | 0±0 | 0±0 | 0±0 | 0±0 |
| GPP + IPP | 11.0±1.4 | 2908.3±361.4 | 109.3±21.6 | 0±0 | 0±0 |
| GPP + IPP (control) | 0±0 | 44.5±3.0 | 63.6±1.7 | 0±0 | 0±0 |
| GPP | 16.8±1.5 | 4040.0±404.1 | 122.2±15.8 | 11.1±1.0 | 0±0 |
| GPP (control) | 0±0 | 40.8±5.1 | 57.8±1.2 | 0±0 | 0±0 |
| FPP + IPP | 0±0 | 0±0 | 0±0 | 1734.9±165.7 | 21.5±3.3 |
| FPP + IPP (control) | 0±0 | 0±0 | 0±0 | 4.0±0.2 | 0±0 |

***Table S8:*** *HMEL037106g1 does not show enzymatic activity with linalool, demonstrating it is not an intermediate in the synthesis of (*E*)-β-ocimene (Fig. S5). Mean amounts (ng) ± standard deviation for each compound across 3 replicates are shown. N=3 for each treatment.*

| | (*E*)-β-Ocimene | (*Z*)-β-Ocimene | Linalool |
|---|---|---|---|
| (*S*)-Linalool | 0±0 | 0±0 | 3182.4±445.5 |
| (*S*)-Linalool (control) | 0±0 | 0±0 | 2698.9±1020.8 |
| (*R*)-Linalool | 0±0 | 0±0 | 3226.6±713 |
| (*R*)-Linalool (control) | 0±0 | 0±0 | 3275.8±350.3 |

***Table S9:** Summary of products from enzymatic assays using precursors from different steps in the pathway (Fig. 1) with both HMEL037106g1 and HMEL037108g1 (Fig. 5A, Table S6, Table S7).*

| Precursors | Enzyme | Products | Activity type |
|---|---|---|---|
| DMAPP + IPP | HMEL037106g1 | Trace (*E*)-β-Ocimene<br>Trace linalool<br>Trace nerolidol | Residual GPS<br>Monoterpene synthase<br>Sesquiterpene synthase |
| DMAPP + IPP | HMEL037108g1 | Trace linalool<br>Trace nerolidol | Residual GPS<br>Monoterpene synthase<br>Sesquiterpene synthase |
| GPP + IPP | HMEL037106g1 | (*E*)-β-Ocimene<br>Trace (Z)-β-Ocimene<br>Linalool | Monoterpene synthase |
| GPP + IPP | HMEL037108g1 | Trace (*E*)-β-Ocimene<br>Linalool | Monoterpene synthase |
| GPP | HMEL037106g1 | (*E*)-β-Ocimene<br>Trace (Z)-β-Ocimene<br>Linalool | Monoterpene synthase |
| GPP | HMEL037108g1 | Trace (*E*)-β-Ocimene<br>Linalool | Monoterpene synthase |
| FPP + IPP | HMEL037106g1 | None | None |
| FPP + IPP | HMEL037108g1 | Nerolidol | Sesquiterpene synthase |

***Table S10:*** *Quantification of experiments characterising IDS activity of HMEL037106g1 (Fig. S7). Only residual IDS activity is detected, with small amounts of (*E*)-β-ocimene, linalool, and nerolidol produced from DMAPP and IPP. No other IDS activity is detected. High amounts of geraniol and farnesol in both experimental and control treatments is due to dephosphorylation of GPP and FPP, respectively. The main function of HMEL037106g1 is the production of (*E*)-β-ocimene from GPP. Mean amounts (ng) ± standard deviation for each compound across 3 replicates are shown. N=3 for each treatment.*

| | (*E*)-β-Ocimene | (*Z*)-β-Ocimene | Linalool | Geraniol | Nerolidol | Farnesol |
|---|---|---|---|---|---|---|
| DMAPP + IPP | 1.7±0.5 | 0±0 | 1±0.1 | 0±0 | 1.6±0.1 | 0±0 |
| DMAPP + IPP (control) | 0±0 | 0±0 | 0±0 | 0±0 | 0±0 | 0±0 |
| GPP + IPP | 325.3±17.4 | 11.3±0.7 | 109.2±10.5 | 1590.1±133.1 | 0±0 | 0±0 |
| GPP + IPP (control) | 2.9±0.2 | 0±0 | 17.6±0.8 | 2300.4±156 | 0±0 | 0±0 |
| FPP + IPP | 0±0 | 0±0 | 0±0 | 0±0 | 15.7±5.1 | 1320.3±114.5 |
| FPP + IPP (control) | 0±0 | 0±0 | 0±0 | 0±0 | 4±0.2 | 1582.0±65.6 |

***Table S11:*** *Quantification of experiments characterising IDS activity of HMEL037108g1 (Fig. S8). TPS activity is again demonstrated by the production of linalool from FPP, and nerolidol from FPP. Only residual IDS activity is detected, by the presence of linalool and nerolidol in treatments with DMAPP and IPP, and nerolidol in the GPP treatment. Geraniol and farnesol are present due to dephosphorylation of remaining GPP and FPP in treatments. Mean amounts (ng) ± standard deviation for each compound across 3 replicates are shown. N=3 for each treatment.*

| | (*E*)-β-Ocimene | Linalool | Geraniol | Nerolidol | Farnesol |
|---|---|---|---|---|---|
| DMAPP + IPP | 0±0 | 3.9±1.6 | 0±0 | 3.5±0.9 | 0±0 |
| DMAPP + IPP (control) | 0±0 | 0±0 | 0±0 | 0±0 | 0±0 |
| GPP + IPP | 12.0±0.8 | 3208.4±261.5 | 290.0±24.5 | 3.6±0.7 | 0±0 |
| GPP + IPP (control) | 0±0 | 30.6±0.7 | 2117.0±184.9 | 0±0 | 0±0 |
| FPP + IPP | 0±0 | 0±0 | 0±0 | 1536.3±61.3 | 983.8±57.5 |
| FPP + IPP (control) | 0±0 | 0±0 | 0±0 | 4.9±5.0 | 1220.8±1105.8 |

***Table S12:*** *Full names of species from Figure 6 in the main text.*

| Abbreviation | Full name |
|---|---|
| *A. gossypii* | *Aphis gossypii* |
| *A. grandis* | *Anthonomus grandis* |
| *A. thaliana* | *Arabidopsis thaliana* |
| *B. mori* | *Bombyx mori* |
| *B. terrestris* | *Bombus terrestris* |
| *C. fumiferana* | *Choristoneura fumiferana* |
| *C. reinhardtii* | *Chlamydomonas reinhardtii* |
| *C. unshiu* | *Citrus reinhardtii* |
| *D. melanogaster* | *Drosophila melanogaster* |
| *D. ponderosae* | *Dendroctonus ponderosae* |
| *F. fujikuroi* | *Fusarium fujikuroi* |
| *G. arboreum* | *Gossypium arboreum* |
| *G. biloba* | *Ginkgo biloba* |
| *H. lupulus* | *Humulus lupulus* |
| *H. melpomene* | *Heliconius melpomene* |
| *H. sapiens* | *Homo sapiens* |
| *I. pini* | *Ips pini* |
| *M. chamomilla* | *Matricaria chamomilla* |
| *M. domestica* | *Malus domestica* |
| *M. histrionica* | *Murgantia histrionica* |
| *M. lewisii* | *Mimulus lewisii* |
| *M. persicae* | *Myzus persicae* |
| *M. piperita* | *Mentha piperita* |
| *N. tabacum* | *Nicotiana tabacum* |
| *N. viridula* | *Nezara viridula* |
| *P. cochleariae* | *Phaedon cochleariae* |
| *P. striolata* | *Phyllotreta striolata* |
| *R. speratus* | *Reticulitermes speratus* |
| *S. cerevisiae* | *Saccharomyces cerevisiae* |
| *T. castaneum* | *Tribolium castaneum* |
